# Design principles for bistable genetic switches emerge from a growth-dependent speed–robustness trade-off

**DOI:** 10.1101/2022.03.31.486579

**Authors:** Kaushik Raj, William T. Z. Wong, Beini Zhang, Radhakrishnan Mahadevan

## Abstract

Bi-stable gene regulatory motifs are found in a wide variety of natural gene regulatory networks and effect transcriptional switching between stable phenotypic states in cells. In synthetic gene regulatory circuits, these architectures can be leveraged to dynamically switch between distinct metabolic states for metabolic engineering and therapeutic applications. However, the lack of modularity and predictability of these motifs in varying environments has limited widespread application, especially since the factors that affect switching characteristics are still unclear. Using a genome-scale metabolic model, we first show that the productivity of a two-stage bioprocess declines as switching is delayed, establishing the value of fast-switching motifs. We then use a mathematical model, along with a newly developed dynamical modeling and continuity analysis framework, to analyze the dynamics and robustness of bi-stable switches over a range of biologically relevant parameter values, and identify a trade-off between the robustness of the motif - the parameter range over which it retains bi-stable function - and the speed at which it effects a phenotypic change. Crucially, by extending our model to incorporate host growth rate through a proteome-partitioning framework, we find that growth rate reshapes the achievable speed-robustness frontier, with the most robust switches attainable at an intermediate growth rate, and that the optimal balance of repressor degradation rates is set by both their relative binding affinities and the growth rate of the host. Using E. coli as a model host, we then constructed a library of 100 transcriptional switches spanning a wide range of switching speeds, experimentally confirming this trade-off and revealing host growth rate as a third design axis - alongside robustness and switching speed - that shapes switch behavior. Together, these results establish design principles for building bi-stable switches with defined switching characteristics, and we anticipate that our library of experimentally validated bi-stable switches will be valuable for effecting phenotypic changes with differing switching-speed requirements in metabolic engineering applications.

## 2 Introduction

Gene regulatory networks in cells confer them the ability to sense their environments and actuate an array of intracellular and extracellular transcription based responses such as the production of substrate assimilation machinery ^1^, initiating cell division ^2^, and fighting extracellular pathogens ^3^. These networks can be decomposed into smaller sub-networks or motifs that are composed of a few parts and carry out specific functions ^4^. One such motif is the bi-stable gene regulatory motif, which helps cells exhibit two distinct stable phenotypic states in response to environmental cues. It is considered to be a primitive form of cellular memory ^5^, since even the transient presence of inducer molecules can effect a stable change in the state of the motif. Such motifs are realized in the native gene regulatory networks of organisms using a double-negative feedback architecture (i.e. two repressors repressing each other’s transcription from inducible promoters) or a positive auto-regulatory architecture (i.e. a transcription factor that induces its own production) ^5^.

Inspired by the design of the bi-stable lambda phage switch, the first synthetic bi-stable motif - the genetic toggle switch Figure 1 was constructed using a pair of repressors (lacI and cI) that inhibit each other’s transcription and is considered to be a pioneering invention in the field of synthetic biology ^6^. Since its conception, the genetic toggle switch, with modifications to its components has been employed in several bio-engineering applications, including in metabolic engineering to decouple growth and production metabolic states to improve chemical production ^7,8^, and in therapeutic applications to detect the presence of inflammation markers ^9,10^. It has also been proposed as a solution for bio-containment of engineered microbes, designed to kill cells that are outside a contained environment^11,12^. The presence of two stable steady states reduces the amount and duration of inducer required to effect a response in the switch, making it a highly valuable induction system in noisy environments to reduce cellular heterogeneity.

**Figure 1:**
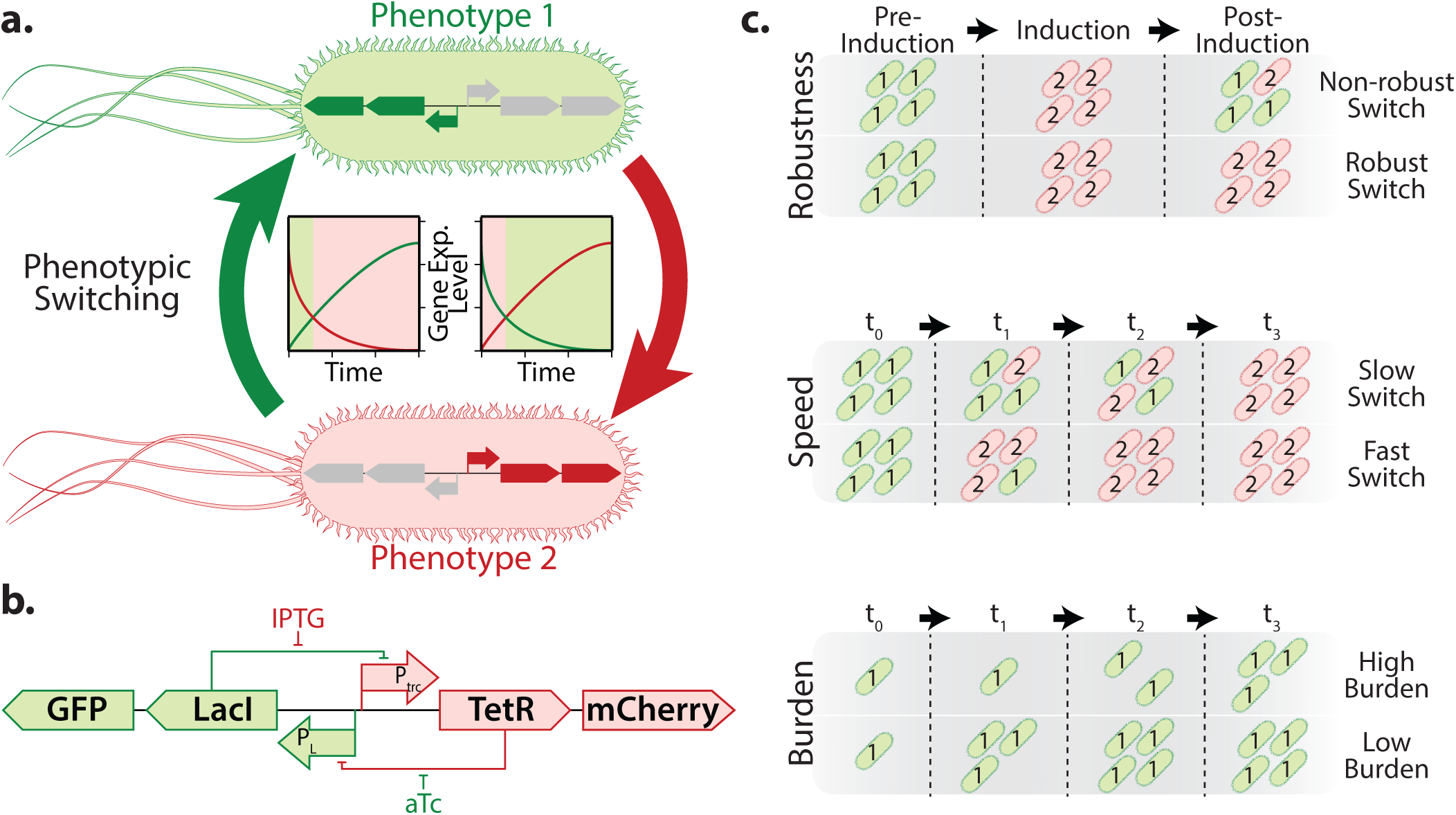
Phenotypic switching in cells and their important functional characteristics. **a.** Gene regulatory networks can be used to effect phenotypic switching in cells to make them express distinct sets of genes or entire pathways in response to stimuli, thereby conferring the cells with distinct phenotypes depending on the stimuli to which they are exposed. **b.** The genetic toggle switch (developed by Litcofsky et al ^13^) is an example of a gene regulatory device which can enable host cells exhibit distinct stable phenotypes through a double negative feedback circuit where two repressors - TetR and LacI which mutually repress each other’s production. We use the genetic toggle switch to study important functional characteristics of switches - illustrated in **c.** Specifically, we are interested in analyzing the effect of different switch constructs on **switch robustness** - the ability of cells to maintain their state even after the input signal is removed, **switch speed** - the speed at which the switch effects the change in phenotypes, and **switch burden** - the impact on cellular growth caused by expression of the switch’s components with a particular emphasis on how that affects the functioning of the switch.

However, despite these proof-of-concept applications, bi-stable motifs have not seen widespread use in scaled up chemical production or therapeutic applications. A major concern with employing this motif for diverse applications, is its context specificity and the lack of predictability of its functional range. Specifically, many versions of the switch have been shown to be bi-stable only in a narrow range of environmental conditions, becoming mono-stable when subject to perturbations in the growth conditions ^6,14^. This precludes their successful deployment as memory elements that require transient induction. Additionally, since these switches use transcription - a relatively slow process in cells, the time taken to switch between the states can be quite large ^7^. This impacts their use in metabolic engineering applications where a fast switch from growth to production stage may be necessary to realize optimal production rates of chemicals ^15^.

Previous studies examining the effects of the growth rates of the host organisms and the metabolic stresses inflicted on them have indicated that there is indeed an interplay between the proper functioning of bi-stable circuits and the conditions in which the host organism is grown ^16–18^. Specifically, a loss of bistability can result when the production of circuit proteins is unable to keep up with their dilution resulting from growth of the host organisms, or when cellular resources are not able to be effectively partitioned to produce the required amounts of circuit proteins due to cellular burden. Consequently, in order to alleviate this burden, there have been efforts to reduce the burden on host cells caused by the expression of the circuit proteins, while maintaining bi-stability ^14^.

The speed of switching between the two stable states is an important characteristic that is often overlooked when designing the toggle switch. Fast acting switches can reduce the latency of engineered biological systems and could reduce inefficiencies in these systems - i.e. the less time spent in transition from one state to another, the less resources wasted in an undesirable metabolic state. This characteristic is highly desirable in microbial chemical production platforms, where a fast switch from a growth to a production phenotype is critical to make such bioprocesses economically feasible ^15^. To quantify the cost of slow switching in such processes, we performed a dynamic flux balance analysis using a genome-scale model of *E. coli* (iML1515) to simulate a typical two-stage microbial chemical production process divided into growth and production stages. In these simulations, we optimized the switch time and the flux distributions of both growth and production stages for bioprocess productivity. Across five representative products, the maximum achievable productivity declined monotonically with increasing switch delay, even when the metabolic strategy was free to re-optimize at each delay (Figure S1). This illustrates concretely why fast switching is desirable for two-stage bioprocesses. Hence, while designing bi-stable motifs for scaled-up platforms, the two characteristics - robustness to cellular resource availability (and consequently varying protein production rates) and the speed of switching are very important to optimize. However, the interplay of robustness and switch speed across the growth regimes encountered in real world applications is yet to be established.

In this work, we first developed a dynamical modeling framework to identify the effects of varying the parameters of a bi-stable motif model on its speed of switching and robustness to varying protein production rates, using the genetic toggle switch as a model circuit. Through simulations on our model using this framework, we observed that the speed of switching in a toggle switch can be improved by increasing the degradation rates of the repressor proteins in the system. However, this improved speed is not without consequence - it results in a decrease in the range of protein production rates over which the toggle switch remains bi-stable, indicating a trade-off between these two characteristics. Extending this analysis to incorporate host growth rate through a proteome-partitioning framework, we found that the achievable trade-off frontier itself varies with growth rate, reflecting the competition between circuit expression and cellular growth. Through a systematic analysis of the switching characteristics over a range of biologically relevant parameter values, we identified optimal parameter ranges that result in maximal robustness of the toggle switch. We then constructed a representative toggle switch with the repressor proteins coupled to a fluorescent reporter to enable live tracking of the switch’s state. Further, we built a large variant library of our newly developed repressor-reporter fusion toggle switch, to validate the speed– robustness trade-off and examine the role of host growth rate in shaping switch behavior that was indicated by our models.

## 3 Results & Discussion

### 3.1 Model description and establishing switch function metrics

The mathematical formulation of the bi-stable architecture that we have proposed (Eq. 1) tracks the concentration of two repressors and consists of 8 parameters that are engineerable *in vivo*. Specifically, the repressor production rates - *k_p_*_1_, *k_p_*_2_ can be altered by modifying the promoter or the ribosome binding sequence to enhance/inhibit transcription and translation rates. The co-operativity of the repressors - *n*_1_, *n*_2_ and their equilibrium dissociation constants - *K_diss_*_1_, *K_diss_*_2_ can be altered by modifying the binding sites of the repressors to each other and to their target promoter sites ^19–21^. The parameters *K_diss_*_1_ and *K_diss_*_2_ also signify the concentration of repressors at which their corresponding promoters result in half-maximal production of the other repressor. Finally, the repressor degradation rates - *k_deg_*_1_, *k_deg_*_2_ can be increased by adding protease affinity tags to the terminii of the proteins. Among these parameters, the repressor protein production and degradation rates are the easiest to design due to the availability of a robust tool to calculate the strength of ribosome binding sites of any DNA sequence ^22^ and genetic parts that result in varying affinities to proteases ^23^. The cooperativity of typically used repressors in synthetic bi-stable motifs - LacI and TetR was taken to be 2 to capture their effective dimeric operator configurations ^24^, following established models of gene regulation described previously ^6,25^. We identified that for bioengineering applications, the speed of switching between the two states and the ability to retain bi-stable behavior in a range of different conditions that impact protein production rates are two important characteristics of a desirable bi-stable motif. Hence, we wished to leverage our newly formulated dynamical modeling framework to examine the effects of varying the two engineerable parameters - protein production and degradation rates on these two functional metrics.

Previous studies have shown that protein production rates in cells are highly variable depending on the growth rate, which itself is governed by an array of environmental conditions to which cells are exposed ^18,26^.

Of the parameters that govern switch function, the repressor production rates are also the most variable once the circuit is designed and deployed. While *K_diss_* and *k_deg_* are set by the choice of binding sites and protease tags and remain largely fixed, the production rates can drift with growth rate, media composition, and the availability of transcriptional and translational resources ^18,26,27^. A switch’s tolerance to variation in its production rates is therefore a meaningful measure of its real world reliability. Accordingly, we define a robustness metric as the switch’s ability to retain bi-stable behavior in response to variations in protein production rates (i.e. the extent of the feasible production rate space over which bistability is preserved - Figure 2a). We note that this usage is distinct from the control-theoretic notion of robustness as the maintenance of stability under parametric uncertainty; here, robustness refers specifically to the size of the bi-stable region in the feasible production rate space. To determine bi-stability regions for a given set of other parameter values, we evaluated the values of the repressor production rates *k_p_*_1_, *k_p_*_2_ that serve as limit points - points that set the threshold for the motif switching from bi-stable to mono-stable behavior, by performing a two-parameter continuity analysis on the toggle switch model. All parameter values within the region bounded by the motif’s limit points will result in a bi-stable configuration, while those outside are mono-stable.

**Figure 2:**
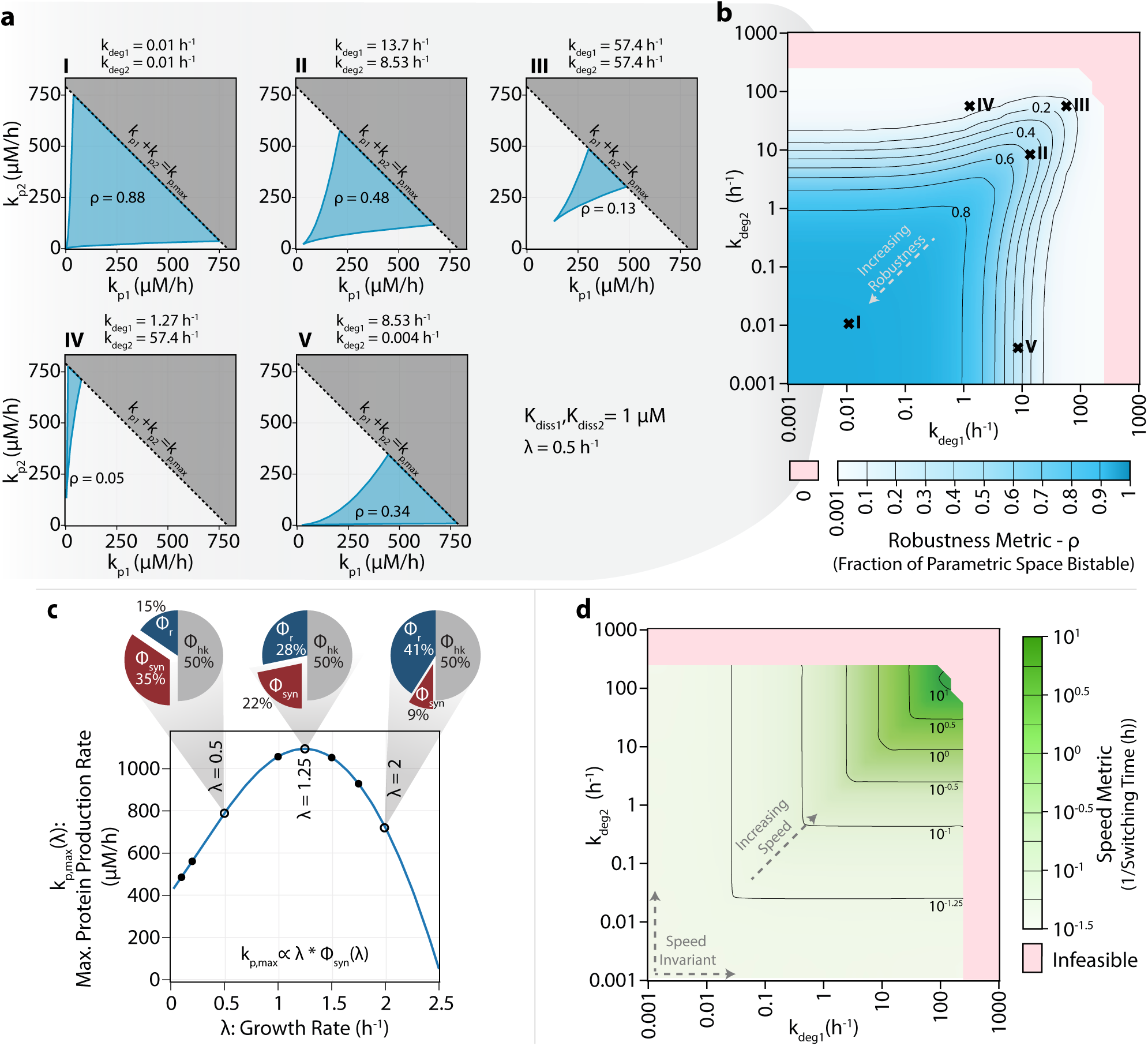
Determination of Robustness & Speed Metrics: **a.** Computation of a robustness metric (*ρ*) at five representative *k_deg_* combinations for a representative set of *K_diss_* and *λ* values. The feasible parameter space for *k_p_* is given by the line *k_p_*_1_ + *k_p_*_2_ *≤ k_p,max_* where *k_p,max_* represents the maximum feasible synthetic protein production rate. The region of bi-stability over the feasible *k_p_* parameter space is computed through continuity analysis and is shown shaded in blue. *ρ* is calculated as the fraction of the feasible parametric space that results in bi-stable configurations. **b.** Robustness landscape over the entire range of experimentally realizable degradation rates at *K_diss_*_1_*, K_diss_*_2_ = 1 and *λ* = 0.5 as representative values. **c.** Determination of maximum feasible synthetic protein production rate (*k_p,max_*) at different growth rates (*λ*). A proteome partition framework is used to divide the steady state proteome in a cell into housekeeping (*ϕ_hk_*), ribosomal (*ϕ_r_*), and synthetic (*ϕ_syn_*) fractions. *k_p,max_* is calculated assuming dilution is only through growth (*λ*). Proteome fractions are illustrated for three representative values of *λ*. **d.** Speed metric landscape over the entire range of feasible repressor degradation rates using the same values of *K_diss_* and *λ* used for b. Speed metric is calculated as the multiplicative inverse of the average switching time from both steady states obtained through kinetic simulations on the toggle switch model.

However, the extent of the feasible production rate space is itself constrained by the growth rate of the host. As the growth rate - *λ* increases, a larger fraction of the proteome must be allocated to ribosomes and growth-associated processes, thereby reducing the budget available for expression of circuit proteins ^28,29^; at the same time, growth dilutes the repressors, contributing to their effective removal alongside active degradation ^27^. In addition to this, the rate of synthetic protein production also varies with the growth rate, in response to overall available transcription-translation machinery ^27^. We therefore incorporate growth rate explicitly into our model, where it enters as - 1. a repressor synthesis rate multiplier (a growth dependent multiplier that affects the realized protein production rate), 2. a dilution term acting on each repressor (i.e. an effective removal rate of *k_deg_* + *λ*), and as 3. a growth-dependent ceiling, *k_p,max_*(*λ*), on the maximum feasible production rate, derived from a proteome-partitioning framework (Figure 2c). A detailed description of the model and assumptions are provided in Methods 5.2 and S1.2. Rather than bounding the production rate space by a fixed maximum, we bound it by the line *k_p_*_1_ + *k_p_*_2_ = *k_p,max_*(*λ*), so that the feasible region reflects the resource constraints imposed at a given growth rate. We computed the robustness metric as the fraction of this feasible production rate space over which the motif remains bi-stable (i.e. robustness metric = 0.1 implies that 10% of all feasible combinations of production rates will result in bi-stable behavior) (Figure 2a). Because *k_p,max_* depends on *λ* (Figure 2c), robustness is intrinsically coupled to the growth rate of the host - a relationship we examine in detail in the following section.

The speed of switching for a given set of parameters can be characterized by calculating the concentration of the two repressors over time upon the addition of inducers, simulated by altering the DNA binding affinities of the corresponding repressors (*K_diss_*_1_, *K_diss_*_2_ set to infinity to simulate the higher affinity of repressors to inducer molecules than their DNA targets) ^30^. In our simulations, the time taken to switch between states was determined as the time taken for either repressor to reach their steady state concentrations and do not deviate further(Figure S2c, d). While the robustness metric directly incorporates all feasible production rates (*k_p_*_1_ and *k_p_*_2_) into its calculation, the speed metric does not, and may vary within the parameter bounds set by our robustness calculations. Hence, we calculated the speed metric as the inverse of the average time taken to switch between the states using repressor production rates at the four extrema of the robustness metric curve (Figure S2a). At a fixed growth rate, we observed that our speed metric is largely intransigent to variations in the repressor production rates, with deviations from the mean value limited to less than 10% for most cases analyzed (Figure S5).

### 3.2 A growth dependent speed-robustness trade-off emerges in the toggle switch model

Having established standard metrics to evaluate the switch’s function for a given parameter set, we first performed a proof-of-concept analysis of our bi-stable motif for a test case where the equilibrium dissociation constants *K_diss_* for both repressors were set to 1 µM and the growth rate *λ* set to 0.5*h^−^*^1^. The value used for *K_diss_* is consistent with *in vivo* observations made in previous studies for the Lac repressor and corresponds to roughly 600 copies of the repressor protein resulting in half-maximal repression of the cognate promoter in a single *E. coli* cell ^31^. We computed robustness values for combinations of degradation rates of the repressor proteins spanning 6 orders of magnitude - ranging from 0.001 *h^−^*^1^ to 1000 *h^−^*^1^. These bounds serve as generous feasibility limits to make our analysis generalizable - hence, though the lower bound corresponds to a protein half-life of 693 hours from degradation alone, we note that in a growing cell the repressors are additionally diluted at the rate of growth, so that the effective removal rate never falls below the growth rate *λ* regardless of the degradation tag used. The robustness metric showed a decreasing trend with increasing degradation rates, with infeasible regions (where no bi-stable configurations were possible) at very high degradation rates corresponding to repressor half-life values <10 seconds (Figure 2b). This is in line with observations made in a previous computational study examining this phenomenon ^32^. Additionally, for this symmetric test case, the maximum robustness was observed when both repressors had the same degradation rate, with robustness values dropping drastically with imbalances in these rates. We observed that imbalances in the degradation rates of the repressors also resulted in impaired switching speeds (Figure 2d). However, in contrast to the trends observed for robustness, the speed of switching increased (corresponding to a decrease in the time required to switch) with increasing degradation rates, reaching a maximum at the highest feasible combination of degradation rates, implying a trade-off between the speed and robustness of the switch. This indicates that switches can be made faster through the addition of protease affinity tags, as suggested previously for a different switch architecture ^33^. Following this, we examined the trends in robustness and switch speeds over the entire range of feasible parameter values. Keeping the co-operativity at 2 and growth rate *λ* = 0.5*h^−^*^1^, we varied the equilibrium dissociation constants of the repressors over 5 orders of magnitude ranging from 0.01 µM to 100 µM (corresponding to 6 copies and 60,000 copies of the repressor protein respectively in each cell, assuming the same volume as *E. coli*). This upper bound on the *K_diss_* values represents approximately 10% of the cell’s entire protein budget ^34^. The trends from our proof-of-concept case, where the switch speed and robustness show opposite correlations to the degradation rates (*k_deg_*) were observed across all *K_diss_* combinations analyzed, indicating that these observations are generalizable (Figure S3, Figure S4).

To better understand the relationship between the two switch metrics, we plotted speed and robustness values for all parameter values from each combination of the dissociation constant *K_diss_* at *λ* = 0.5*h^−^*^1^ (Figure 3a, Figure S6a, Figure S7). In general, as speed increases, both the maximum robustness and the range of robustness values achievable decline - bolstering the trade-off we alluded to earlier. Further, in each of these plots, we traced the Pareto fronts formed by the robustness and speed metrics, which show a clear trade-off between them. Interestingly, at the points constituting the Pareto front in each of these plots, we find that the ratio of the *effective* degradation rates, which include the contribution of growth-mediated dilution: *r* = (*k_deg_*_1_ + *λ*)*/*(*k_deg_*_2_ + *λ*) was roughly equal to the ratio of the dissociation constants - *K_diss_*_2_*/K_diss_*_1_. This relationship has an intuitive interpretation - *K_diss_*_2_ reflects how effectively repressor 2 represses the production of repressor 1, so when repressor 2 is a weak repressor (large *K_diss_*_2_), a correspondingly high degradation rate of repressor 1 is required to restore balance. As the dissociation constants become more asymmetric, the optimal degradation imbalance grows correspondingly, seemingly restoring the balance disrupted by the differing binding affinities of the two repressors to their cognate promoter targets. This finding is important when designing bi-stable motifs with two different repressors, as in the genetic toggle switch, in order to determine the optimal protease tag to add to each repressor. For the symmetric test case with equal binding affinities (Figure 3a), this reduces to balanced degradation (*r_opt_* = 1).

**Figure 3:**
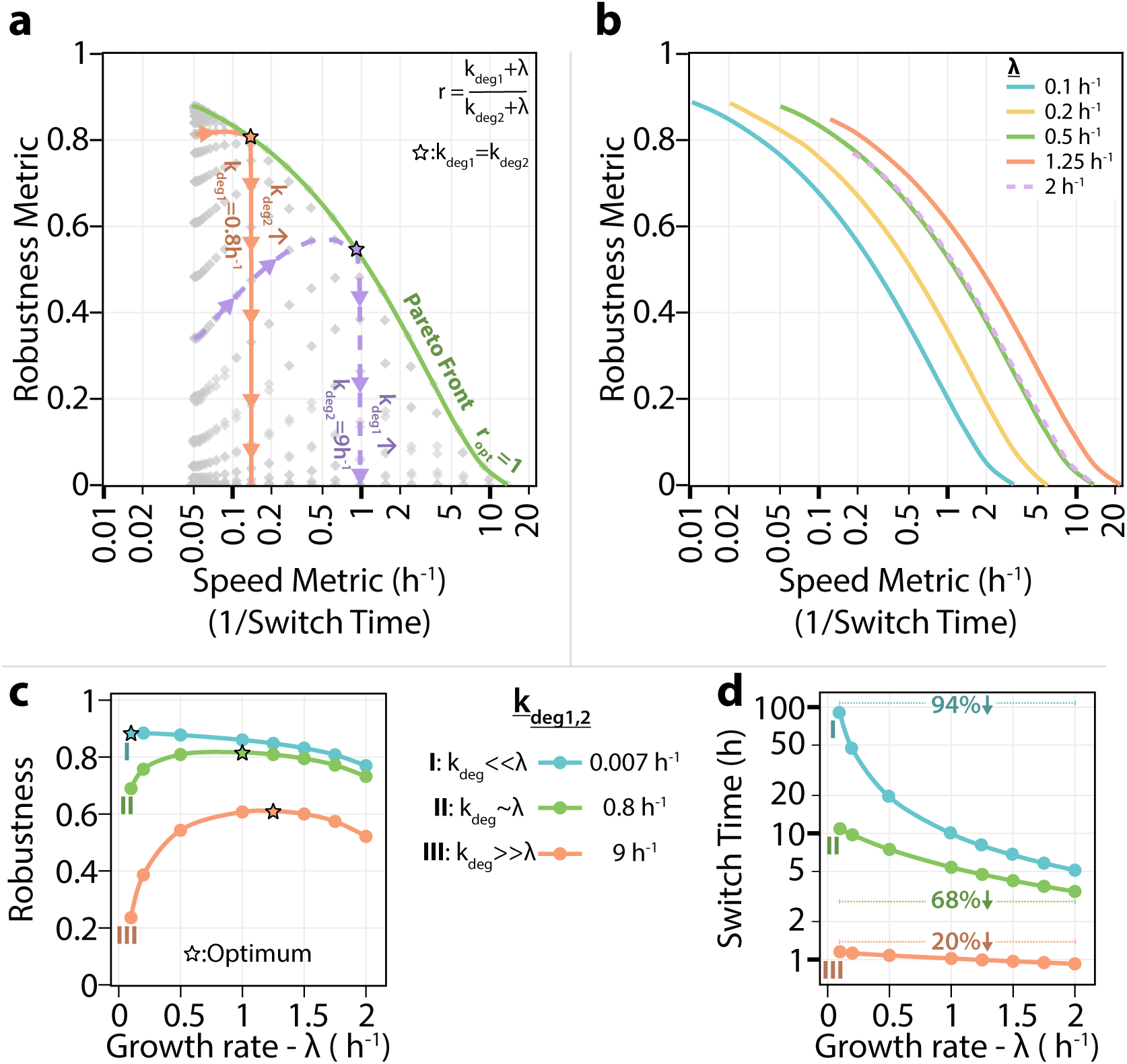
Growth rate shapes a speed-robustness trade-off and defines optimal switch design. **a.** Speed and robustness metrics for all degradation-rate combinations at *K_diss_*_1_ = *K_diss_*_2_ = 1 µM, with the Pareto front in green. The front is traced when the effective degradation rates are balanced, i.e. the ratio *r* = (*k_deg_*_1_ + *λ*)*/*(*k_deg_*_2_ + *λ*) equals its optimal value *r_opt_* = 1 (star: *k_deg_*_1_ = *k_deg_*_2_). Two trajectories show the effect of tuning one repressor at a time: varying *k_deg_*_2_ at fixed *k_deg_*_1_ = 0.8 h*^−^*^1^ (orange) and varying *k_deg_*_1_ at fixed *k_deg_*_2_ = 9 h*^−^*^1^ (purple). **b.** Pareto fronts at five growth rates, showing that the achievable frontier itself shifts with growth rate. **c.** Robustness versus growth rate at three representative degradation rates spanning three regimes: (I) *k_deg_ ≪ λ* (dilution-dominated), (II) *k_deg_ ∼ λ* (mixed), and (III) *k_deg_ ≫ λ* (degradation-dominated); stars mark the growth rate of maximal robustness. **d.** Switch time versus growth rate for the same three regimes, with the reduction across the growth-rate range indicated.

It should be noted that this optimal ratio is realizable only where the degradation rates are within the feasible parameter grid of *k_deg_* values we have selected for simulation; outside this range, the frontier is instead bounded by the feasible limits on the degradation rates themselves (indicated by dashed Pareto front in Figure S6a, Figure S7). In addition, examining the two single-repressor trajectories in Figure 3a and Figure S6a revealed that switching speed is governed by the slower of the two effective degradation rates: increasing only one degradation rate accelerates switching until the two rates equalize, beyond which speed saturates while robustness continues to vary. Efficient traversal of the speed-robustness frontier therefore requires scaling both degradation rates together while maintaining an optimal ratio, rather than tuning either repressor alone.

Because growth rate enters both the dilution of the repressors and, through proteome partitioning, the maximum feasible production rate, it reshapes the achievable speed-robustness frontier (Figure 3b, Figure S6b, Figure S8). We examined the nature of this effect in further detail for two different cases - when *K_diss_*_1_=*K_diss_*_2_ = 1*µM* and when *K_diss_*_1_ = 0.01*µM, K_diss_*_2_ = 1*µM* . This revealed that the effect of growth rate depends on the regime of degradation rates relative to the growth rate (Figure 3c, d, Figure S6c, d). When degradation is negligible relative to growth (*k_deg_ ≪ λ*), switch metrics are governed almost entirely by dilution: switching accelerates strongly with growth rate, and robustness declines monotonically as growth increases. When degradation dominates (*k_deg_ ≫ λ*), switch speed becomes nearly independent of growth rate, and the growth-dependence of robustness instead reflects changes in the effective production rate, reaching a maximum at *λ* = 1.25*h^−^*^1^, where the growth-dependent production budget is largest (Figure 2c, Figure 3c, Figure S6c). At intermediate degradation (*k_deg_ ∼ λ*), both effects contribute and robustness peaks near *λ* = 1 *h^−^*^1^. Consistent with speed being governed by degradation, the reduction in switch time with increasing growth rate was largest when degradation was slow (a 94% reduction across the growth-rate range for *k_deg_ ≪ λ*) and progressively smaller as degradation increased (68% and 20% for the intermediate and high-degradation regimes; Figure 3d, Figure S6d). In all regimes, however, the underlying speed-robustness trade-off is preserved.

Finally, we observed that the robustness, and consequently the range of feasible degradation rates, became smaller with increasing *K_diss_* (representing impaired repression). These results indicate that a bi-stable motif employing repressors with enhanced DNA binding ability will result in a highly robust construct, allowing better tuning of the switch speed through the addition of protease affinity tags than similar motifs that use weakly binding repressors or those with leaky expression. While the robustness metric showed significant differences with changing *K_diss_* values, the speed metric remained constant over the entire range of *K_diss_* values at a given growth rate, with only the feasibility region becoming smaller with poorer repressor-DNA binding (Figure S4). Along with our earlier observation that switch speed does not change with repressor production rates, this indicates that, at a fixed growth rate, the degradation rate of the repressor proteins is the primary determinant of switch speed, further demonstrating the importance of using protease affinity tags to accelerate bi-stable motifs.

### 3.3 Repressor-reporter fusion to characterize switch state in real time

Following the discovery of design guidelines to build better bi-stable motifs, we wished to validate our model findings by building faster switching variants of the genetic toggle switch described by Litcofsky et al ^13^. Additionally, we also wanted to demonstrate the presence of a speed-robustness trade-off experimentally. In order to monitor the speed of switching, it is necessary to track the concentrations of the repressor proteins *in vivo* to determine when a state change has occurred. However, in the original design of the toggle switch, the repressor and fluorescent reporter proteins are de-coupled, precluding a quantitative analysis of repressor concentrations by simply measuring the fluorescence intensities of the reporters. Hence, we aimed to construct a fusion variant of the genetic toggle switch where the repressor and reporter proteins are physically coupled to each other by means of the flexible protein linker - (GGGGS)_2_ ^35^. In order to determine the optimal configuration of the repressor-reporter fusions (N-terminal vs C-terminal fusions) that resulted in similar repression characteristics as their non-fusion counterparts, we constructed all possible variants of fusion constructs of the LacI repressor with the fluorescent protein GFP, and the TetR repressor with the fluorescent protein mCherry. We then inserted these constructs into test circuits where they would repress expression of a fluorescent protein and can be induced with their cognate inducer (IPTG for LacI and aTc for TetR). The test circuits used for each repressor, shown in Figure 4 consist of GFP or mCherry under the control of either the *p_L_* or *pTrc* promoter respectively. The fusion constructs with TetR or LacI were expressed constitutively as required in the test plasmid constructs, repressing production of the fluorescent reporters GFP and mCherry, until they are sequestered through the addition of aTC or IPTG respectively..

In our designs, we observed that neither configuration for both repressors resulted in a significant increase in leaky expression (expression of downstream genes without the addition of inducers) from their target promoters, indicating that the fusion constructs are at least as good as their wild-type counterparts in DNA binding ability (Figure S9). Additionally, we also examined the ability of the repressors to be induced by their corresponding inducer molecules. In all cases, the fusion constructs showed weaker induction characteristics than their wild-type counterparts, with higher concentrations of inducer being required for an equivalent de-repression of their promoters. We chose the fusion protein configuration that had a lower *K_M_* (inducer concentration required for half maximal expression) and lower variance in induction characteristics. Hence, the TetR-mCherry (TetR at N-terminus and mCherry at C-terminus) and GFP-LacI(GFP at N-terminus and LacI at C-terminus) fusion proteins were chosen as optimal constructs for further experiments.

**Figure 4:**
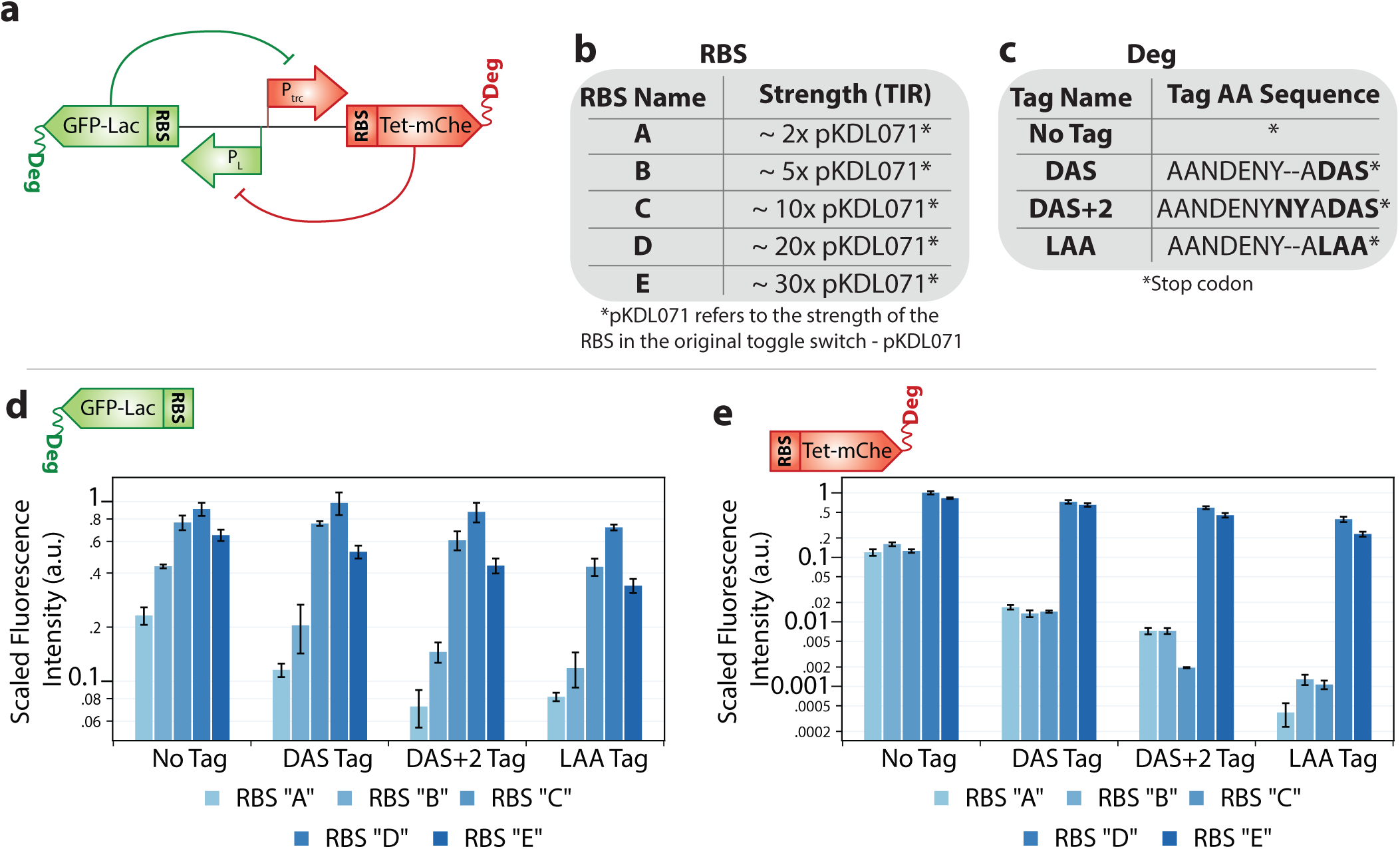
Fusion Toggle Switch Study Speed-Robustness Trade-off. **a.** An illustration of the fusion toggle switch with GFP-LacI and TetR-mCherry with modified ribosome binding site (RBS) and added degradation tags. **b.** The five RBS strengths (as predicted by RBS calculator v2.1^22^) chosen for each repressor relative to its value on the original pKDL071 toggle switch designed by Litcofsky et al ^13^. The queries and translation rates corresponding to each RBS level is shown in Table S2. **c.** Degradation tags and their amino acid sequences added to the repressors for toggle switch library construction. Modified versions of the SsrA degradation tag that show different degradation rates were chosen. **d.** GFP-LacI and **e.** TetR-mCherry fluorescence intensities determined using a microplate reader (n*≥*3) for each of the 20 RBS-degradation tag combinatorial variants of repressor-reporter fusion proteins, organized by the degradation tag used. Refer to Methods 5.5 for experimental details.

### 3.4 Constructing and characterizing a diverse toggle switch library

Guided by these design principles, we set out to construct a library of toggle switch variants spanning a range of repressor production and degradation rates, with the goal of obtaining switches that are both fast-switching and bi-stable. We began by assembling a single representative fusion toggle switch using GFP-LacI and TetR-mCherry as the repressors, with the ribosome binding strength of either repressor engineered to mimic the values found in the original pKDL071 version of the toggle switch ^13^ (Table S2). This construct expressed the repressors as intended, displaying high mCherry fluorescence (and correspondingly high TetR concentrations) when grown on IPTG, and high GFP fluorescence (and correspondingly high LacI concentrations) when grown on aTc (Figure S10). It was, however, mono-stable (Supplementary data file: SI_Data), reverting to a TetR-high state after the removal of aTc, consistent with the calculated RBS values placing the construct’s production rates outside the bi-stable region of the parameter space. Rather than a setback, this provided a direct test of our model’s central prediction - that bi-stability can be restored by rebalancing the repressor production and degradation rates. We therefore used this construct as the scaffold from which to build our variant library, systematically varying the ribosome binding strengths and degradation tags of both repressors to bring the switch into its bi-stable regime.

Next, we designed variants of the fusion toggle switch with varying ribosome binding site(RBS) strengths and protease affinity tags (equivalent to varying *k_p_* and *k_deg_* values respectively in the model) to create constructs that result in a range of protein production and degradation rates, as required to validate our model findings. To this end, we designed RBSs that we expected to show between 2 and 30 times the strength of the RBS in the original toggle switch - pKDL071 using the RBS calculator (v2.1) ^22^ (Table S2). In order to vary the degradation rates of the toggle switch proteins, we leveraged the native *ClpXP* protease in *E. coli* by tagging the repressors with the SsrA tag and its variants that have been used in previous studies to show graded degradation rates ^23,36^. A list of RBS strengths and degradation tags we chose for each repressor are shown in Figure 4. This resulted in 20 variants of each fusion repressor (5 RBS levels * 4 degradation levels), composed of all possible combinations of the RBSs and degradation tags.

To shortlist functioning RBS and degradation tags for our final combinatorial library, we first examined the protein production and degradation rates of the 20 variants of each fusion repressor leaving the other repressor unchanged (40 constructs in total), by transforming plasmids harboring these constructs into *E. coli* MG1655 (ΔlacI) ^14^ and inducing their expression with IPTG or aTc. We then monitored the levels of repressor-reporter fusion proteins produced by the RBS-degradation tag combinations by recording the fluorescence intensity of the reporter in a microplate reader. As expected, the addition of degradation tags resulted in progressively higher rates of degradation of the repressors and consequently, lower steady state fluorescence intensities from the reporter attached to them (Figure 4d, e and Figure S11). Furthermore, the strengths of the degradation tags indicated by these results are largely in line with observations made in previous studies (‘LAA’ has the highest strength, followed by ‘DAS+2’ and then ‘DAS’) ^36^. The relative rates of degradation, as indicated by the reduction in the steady-state fluorescence intensity at each RBS level were estimated and have been listed in Table S3 and Table S4.

However, RBS strengths did not conform with our design parameters, specifically for TetR-mCherry, with many RBSs constructs showing minimal variation (Figure 4d, e and Figure S11). Hence, we shortlisted 10 combinations of RBS-degradation tag variants for each repressor, that showed the largest variability in the fluorescence intensities (and consequently the repressor production/degradation rates) in order to obtain a highly variable combinatorial library of toggle switch variants and thereby examine a large portion of the design space. We then constructed plasmids containing all possible combinations of these shortlisted variants using the ligase cycling reaction method ^37^, sequence confirmed the 100 resulting plasmids containing toggle switch variants, and transformed them into *E. coli* MG1655 (ΔlacI) for final characterization. As we will see in following sections, this library contained a large number of constructs that are bi-stable across repeated passaging without inducers (Figure 6b,c), confirming that rebalancing the production and degradation rates restores bi-stability as predicted.

### 3.5 Switch variants show a wide range of switching speeds

Leveraging our previously described high-throughput phenotyping platform ^38^, we examined the switching speed of our 100 switch variants in triplicate. We monitored the time taken by either repressor to reach its steady state maximum and minimum upon the addition of the corresponding inducer (Figure 5a). As expected, adding degradation tags and thereby increasing the degradation rates of the Tet repressor resulted in a generally faster rate of the Tet repressor reaching its steady state value in both states. In particular, the addition of degradation tags greatly reduced the time taken to degrade TetR from cells that are switching into the Lac high state, as indicated by the large reduction in the Tet degradation time for our library variants compared to the original construct (Figure 5b). We observed a similar trend with the Lac repressor as well, with many library variants with degradation tags showing a shorter time to reach the steady state value (Figure 5c). There were however, some Lac repressors with relatively low degradation rates (as predicted in Table S3 and Table S4) that reached steady state much faster than those with higher degradation rates, possibly due to their extremely low RBS levels, resulting in very small steady state values.

**Figure 5:**
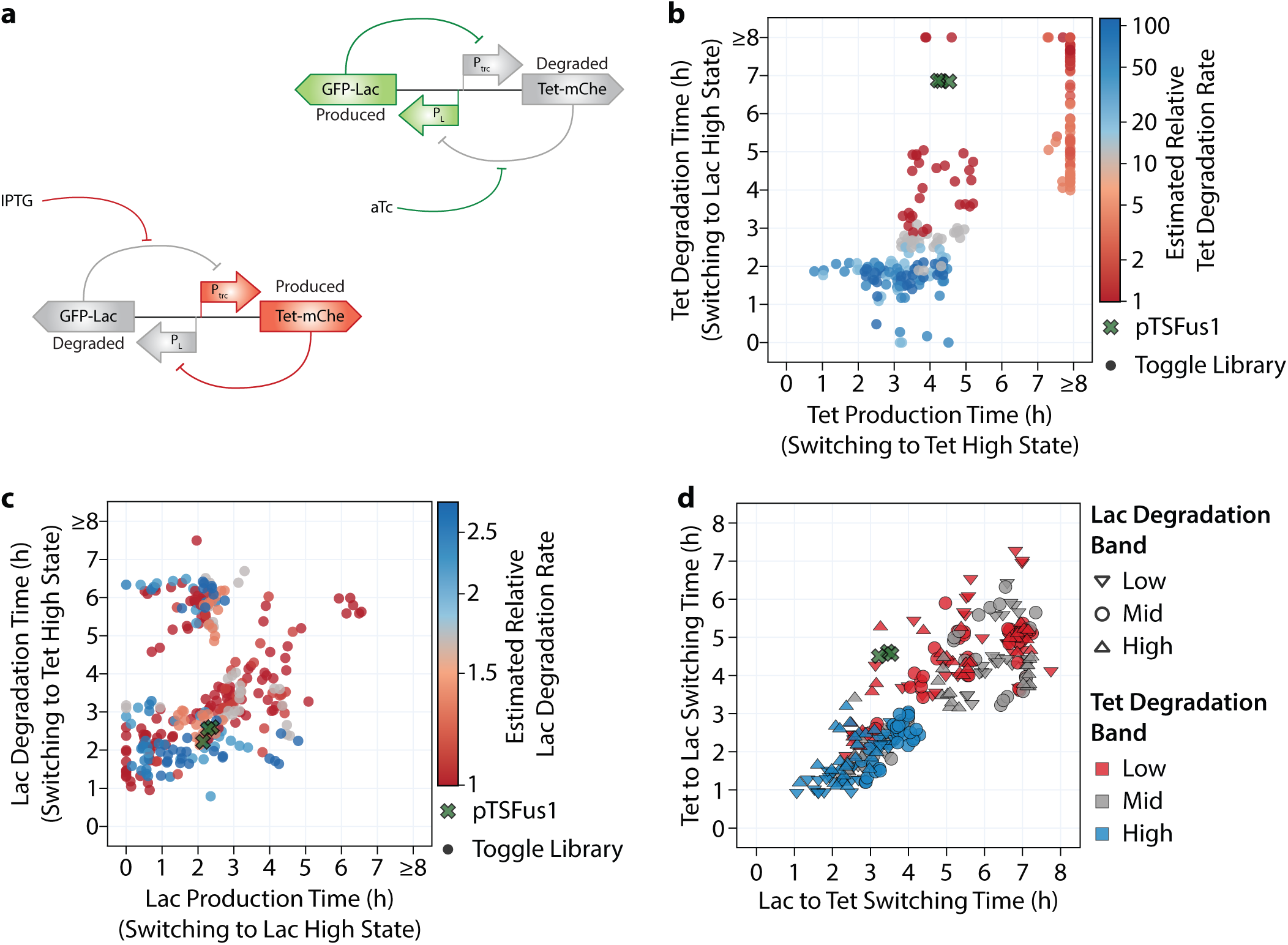
Switching time vs Degradation rates. **a.** Illustration showing the addition of different inducers and which repressors are produced or degraded in each case. The time taken by either repressor to be produced and degraded in the appropriate state is calculated. **b.** Effect of degradation tags on the Tet repressor in reaching steady state concentrations. **c.** Effect of degradation tags on the Lac repressor in reaching steady state concentrations. **d.** Effect of degradation tags on the switching time (calculated as the average of the time taken to produce and degrade the appropriate repressor for each state), with strains classified based on degradation levels estimated in Table S3 and Table S4. Refer to Methods 5.5 for experimental details.

**Figure 6:**
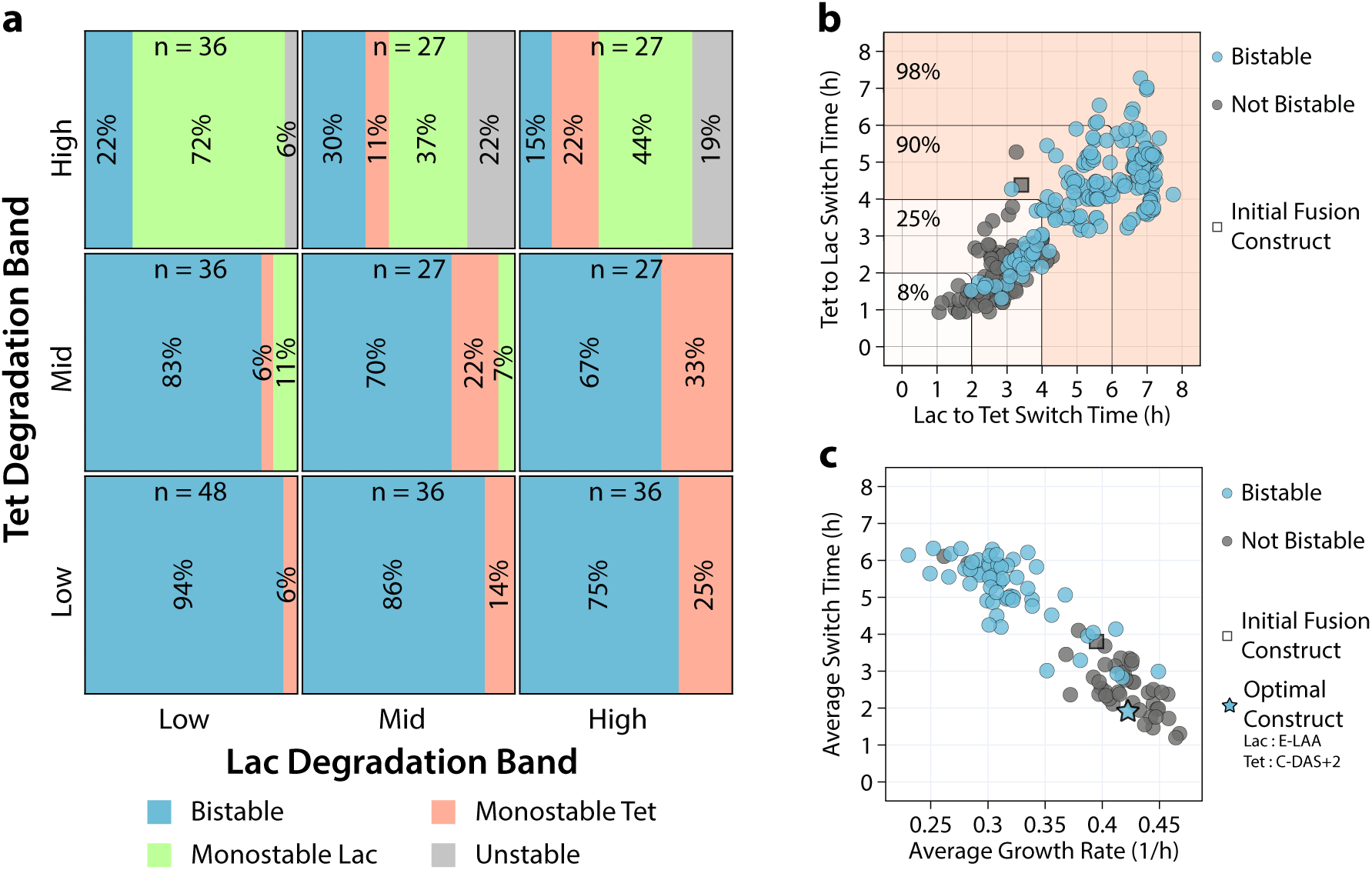
Trade-offs in toggle switch behavior: **a.** Effect of varying degradation levels of the Lac and Tet repressor on the bi-stability of individual replicates. The percent of bi-stable, mono-stable and unstable constructs are shown within the boxes representing each degradation level combination. Constructs were classified into different degradation levels as shown in Table S3 and Table S4. Overall data for each construct is shown in Figure S14. **b.** The Lac state and Tet state switching times for individual replicates, classified based on stability. The percent of bi-stable replicates at each level of Tet and Lac switching times is shown within the plot. Overall data for each construct is shown in Figure S15. **c.** A trade-off between the average switching time (calculated as the average of the two state-switching times) and the growth rate of the each strain, classified based on stability. Constructs were designated bi-stable if all individual replicates were bi-stable. Individual replicate data shown in Figure S16. Refer to Methods 5.5 for experimental details.

Nevertheless, upon comparing the effect of various degradation rates on the overall switching time (mean value of the time taken to degrade one repressor and produce another), we found that those constructs with higher degradation rates, generally resulted in faster switching to either state (Figure 5d) as predicted by our model. Specifically, we found that the degradation tags on the Tet repressor are highly effective in mediating a faster switch, likely due to the higher relative degradation rates in the Tet repressor than the Lac repressor (Table S3 and Table S4). However, several constructs with intermediate degradation rates resulted in very slow switching. Upon a closer analysis, we found that these were constructs with extremely high protein production rates, which resulted in very slow growth and consequently, slower switching than even those constructs without degradation tags. This can be clearly seen when the switching times are plotted against the growth rates of the corresponding cells, where a number of variants with intermediate degradation rates result in very slow growth and therefore slow switching (Figure S12). A principal component analysis performed on these metrics also confirmed this observation, showing a strong negative correlation between the time taken to degrade repressor proteins and the host strain’s growth rate for either state(Figure S13c). In other words, constructs with intermediate degradation rates resulted in slow switching only when the growth rate was low (Figure S13a, c). Hence, the growth rate of the resultant strains is an important parameter that also needs to be considered while designing switches, further bolstering our model predictions. Regardless, our variant library consists of switches that show a wide range of switching time, many of which are faster than the initial construct, indicating that varying the degradation rates is indeed an effective strategy to alter switching speeds. Furthermore, we generally observed that higher degradation rates result in faster switching when the growth rate is constant (Figure S12 and Figure S13).

### 3.6 Trade-offs in switch function - experimental validation

Having confirmed the effect of adding degradation tags on the speed of switching, we examined the ability of our switch variants to retain their state after removal of their corresponding inducer i.e. the ability to maintain high levels of Tet after IPTG is removed and maintain high levels of Lac after aTc is removed. Based on their ability to retain their state after three consecutive transfers with no inducer, we classified the individual experimental replicates as - Bi-stable when both states are stable, Mono-stable when only either one of the Tet or Lac state is stable and unstable when the construct is unable to retain either state (Figure 6a). Our results show that a majority of strains with slow degrading repressors are bi-stable, with the proportion of bi-stable strains decreasing as degradation rates are increased. We find the lowest proportion of bi-stable strains in the constructs with the highest degradation rates on both repressors. This is in remarkable agreement with our model findings where robustness (i.e. the region of bi-stability and consequently the number of bi-stable constructs) decreased with increasing degradation rates. Additionally, we also found that increasing the degradation rate of one repressor resulted in more constructs becoming mono-stable towards the other. Additionally, imbalanced degradation rates also resulted in lower robustness, as predicted by our model. We observed the same trends when the experimental replicates of each strain were combined to obtain the overall bi-stability of a construct as opposed to the individual replicates (Figure S14).

Next, we compared the number of bi-stable strains at each level of switching speed to study any possible correlations between the robustness and the speed metric. Once again, in remarkable agreement with our model predictions, we found a monotonically decreasing trend in the proportion of bi-stable constructs, with increasing switching speeds (or decreasing switching times) (Figure 6b and Figure S15). This confirms the presence of a trade-off between the switching speed and robustness in the genetic toggle switch, indicating the need to carefully design switches with appropriate protein production and degradation rates. Importantly, we identify a range of different switch constructs that are fast switching while still being bi-stable. Further, we also observed that there was an inverse relationship between the average switching time and growth rates among our experimental constructs, in agreement with model predictions. Additionally, since we uncovered previously that the growth rate of the host varies significantly depending on the switch configuration and fast growth is a highly desirable characteristic in metabolic engineering applications, we wished to identify optimally performing switch constructs that switch fast and remain bi-stable, while not imparting a severe growth burden on the host. Comparing the switching time, growth rate and the bi-stability of each strain, we found that all replicates of the construct with the ‘E-LAA’ Lac repressor and the ‘C-DAS+2’ Tet repressor resulted in optimal characteristics and can be investigated for further improvement (Figure 6c and Figure S16).

## 4 Conclusions

Gene regulatory circuits enable highly effective control of cellular states in engineered bacteria by modifying environmental conditions or through the addition of exogenous chemical signals. This allows engineered cells to act as microscopic controllers - sensing their environments and actuating an appropriate genetic response. Such systems show immense potential for use in metabolic engineering and therapeutic applications. Specifically, bi-stable motifs can greatly reduce variability in cellular phenotypes by forcing cells to exist in one of two stable steady states. However, the design of such systems has been a difficult endeavour in the past due to an incomplete understanding of how each part of the circuit affects its function.

In this work, we have shown that the design choices made during the construction of bi-stable motifs impact their function to a great extent. Using a mass-action kinetics based model of the toggle switch, we characterized a trade-off between the speed and robustness of bi-stable motifs, and identified ways to engineer the toggle switch to accelerate switching between the states. This trade-off has direct consequences for metabolic engineering: the time at which microbes are switched from growth to production phenotypes, and the duration of the switch, play an important role in determining the economic feasibility of a bio-process. A slow switch to the production stage could result in impaired chemical production and thereby render such processes economically infeasible; indeed, our dynamic flux balance analysis of a two-stage bioprocess showed that the achievable productivity declines monotonically with increasing switch delay across a range of products.

While fast-acting switches are highly desirable for metabolic engineering and other applications, this speed comes at the cost of robustness to variation in protein production rates. This trade-off places a limit on how much the speed of switching can be improved before the motif loses bi-stability, and conversely suggests that slow-acting switches are preferable for applications where high robustness is required. Critically, by extending our model to incorporate host growth rate through a proteome-partitioning framework, we found that growth rate is itself a determinant of switch behavior: it reshapes the achievable speed-robustness frontier, with the most robust switches attainable at an intermediate growth rate where the proteome budget available for circuit expression is largest. This analysis also yielded a concrete design rule - the most robust switch at any given speed is achieved when the effective degradation rates of the two repressors, including growth-mediated dilution, are balanced in proportion to their binding affinities, i.e. (*k_deg_*_1_ + *λ*)*/*(*k_deg_*_2_ + *λ*) = *K_diss_*_2_*/K_diss_*_1_. Switch speed can then be tuned by scaling both degradation rates together along this optimal ratio, whereas tuning either repressor alone sacrifices robustness without a commensurate gain in speed.

Our model and experiments show that robustness, switching speed, and host cell growth rate are mutually constrained. Switches employing higher degradation rates can be made bi-stable by increasing the production rates of their repressors, but this imposes a growth burden on the host that in turn slows switching; conversely, strains that grow extremely fast also tend towards unstable bi-stable motifs. These three quantities are therefore at odds with each other, and the key to constructing optimal switches lies in carefully negotiating the effects of the switch’s components on all three. Taken together, these results provide a set of design principles for bi-stable switches: match the effective degradation ratio to the repressors’ binding-affinity ratio for maximal robustness, scale both degradation rates together to traverse the speed-robustness trade-off, and account for the host growth rate, which sets both the accessible frontier and the optimal operating point. We believe these findings will be influential in designing bi-stable motifs for metabolic engineering and therapeutic applications. Moreover, our generalized dynamical modeling framework can be used to analyze other regulatory motifs and could be a valuable tool for designing complex biological circuits. Finally, we constructed a modified fusion variant of the toggle switch in which the repressor proteins carry fluorescent reporters, which could serve as a valuable tool in dynamic control applications, since the state of the switch can be read out non-invasively by simply monitoring cellular fluorescence.

In this study, several constructs in the highest degradation band were not bi-stable or mono-stable in either state (Figure 6a). This deviates from our predictions since, based on our deterministic model, toggle switch constructs can only be bi-stable or mono-stable in one state. Stochastic effects, in which a change in switch state is effected by noise in the system, could explain this and should be considered in future modeling efforts. Furthermore, the interplay between the growth rate of host cells and circuit function should be explored further using models of cellular resource allocation that predict growth rate as an emergent consequence of heterologous protein expression, rather than as an imposed parameter as in the present work. One potential way to alleviate the burden of expressing the toggle switch proteins is to build a variant library in a single-copy vector or through genomic integration of the circuit, as shown in a recent study ^14^.

Nevertheless, our combinatorial library of toggle switch variants shows a wide range of switching speeds while remaining bi-stable, and could be used to enhance the production of a diverse range of target compounds by decoupling growth and production phenotypes. We anticipate that this work will encourage the widespread use of bi-stable motifs in bioengineering for the dynamic control of metabolism.

## 5 Methods

### 5.1 Mathematical model of the growth-coupled toggle switch

The genetic toggle switch was chosen as the model circuit to study the effect of bi-stable motifs. It consists of two repressors that inhibit each others’ transcription, resulting in a net positive feedback motif (Figure 1). The state of the switch can be altered by means of external inducers, which act by binding to and inactivating their corresponding repressor targets. The proposed mathematical model for the genetic toggle switch (shown in Eq. 1) was generated using mass action kinetics where *Repressor*1 and *Repressor*2 represent the repressors LacI and TetR in the case of the genetic toggle switch. For the sake of simplicity, we assumed that repressors binding to their cognate promoter targets occurred at a much faster time-scale than the production of the repressor proteins, thereby resulting in an equilibrium between the repressors and their targets. To simulate the effect of the addition of inducer molecules, the ability of the repressor species to bind to their cognate promoter targets was impacted by increasing the values of the parameter *K_diss_* to a very high value, where the repressor could no longer inhibit gene expression from its promoter target.

To capture the dependence of switch function on the physiological state of the host, the model incorporates the growth rate *λ* in two ways. First, growth dilutes the repressor proteins, contributing to their effective removal alongside active degradation; the removal rate of each repressor is therefore *k_deg_* + *λ*. Second, the rate of protein synthesis depends on growth rate: we represent the production rate of each repressor by a lumped parameter *k_p_*, distinct for each repressor and constant over the course of a simulation, and relate the *effective* production rate *k_p,eff_* to a *designed* production rate *k_p_* through a growth-dependent synthesis multiplier, kp,eff = kp (1 − e−λ/λ∗), following Hintsche et al ^27^. Here *k*_p_ is the production rate that would be specified by the choice of promoter and ribosome binding site, while *k_p,eff_* is the rate realized at a given growth rate; the multiplier captures the empirical reduction in expression capacity as growth slows, with *λ^∗^* = 0.24 *h^−^*^1^. The resulting model is given in Eq. 1.

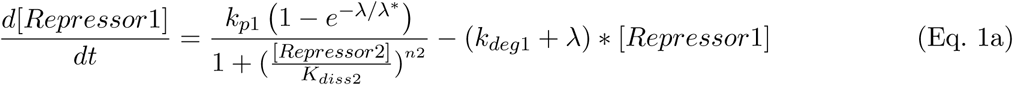

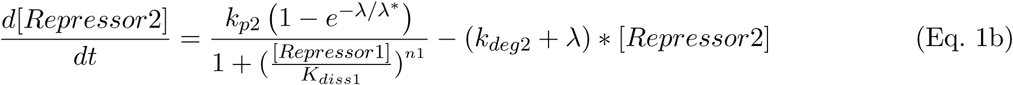

where

[*Repressor*1], [*Repressor*2]: Concentration of repressors (*µM*)
*k_p_*_1_*, k_p_*_2_: Designed production rate of repressors (*µM/h*) *λ*: Specific growth rate of the host (*h^−^*^1^)
*λ^∗^*: Growth rate constant of the synthesis multiplier (*h^−^*^1^)
*K_diss_*_1_*, K_diss_*_2_: Equilibrium dissociation constant for repressors binding to cognate promoters (*µM*)
*n*_1_*, n*_2_: Cooperativity of repressor binding
*k_deg_*_1_*, k_deg_*_2_: Degradation rate of repressors (*h^−^*^1^)

### 5.2 Growth-dependent constraint on protein production

The maximum rate at which a host cell can produce the repressor proteins is limited by the fraction of its proteome that can be allocated to their expression. We estimated this limit using a proteome-partitioning framework ^28,29^. The proteome is divided into a growth-independent housekeeping fraction *ϕ_hk_* - the fraction devoted to core cellular processes whose expression does not vary with growth rate which accounts for approximately 50% of total protein, a ribosomal (and growth-associated) fraction *ϕ_r_* that increases linearly with growth rate as *ϕ_r_*(*λ*) = *ϕ_r_*_0_ + *w_r_λ* - where *ϕ_r_*_0_ is its growth-rate-independent intercept and *w_r_* is its slope with respect to growth rate, and the remaining fraction available for expression of the circuit proteins, *ϕ_syn_*(*λ*) = 1 *−ϕ_hk_ −ϕ_r_*(*λ*). As growth rate increases, the expansion of the ribosomal fraction reduces *ϕ_syn_*, tightening the budget available for circuit expression.

We emphasize that *ϕ_syn_*(*λ*) represents a maximum feasibility limit on the synthetic proteome fraction rather than a realistic allocation. We account only for the two largest obligatory sectors - the housekeeping and ribosomal fractions and assign the entire remainder to synthetic protein. Other proteome sectors that would in practice further reduce the available fraction, such as those devoted to carbon catabolism, transport, and metabolic enzymes, are not included. The resulting ceiling should therefore be interpreted as an upper bound on the feasible production rate, set by the bare minimum proteome commitments required for growth.

The maximum feasible steady-state concentration of synthetic protein at a given growth rate is then *P_syn,max_*(*λ*) = *P_tot_ ϕ_syn_*(*λ*), where *P_tot_* is the total protein concentration in exponentially growing *E. coli*. At steady state, the production of a protein is balanced by its removal; taking growth-mediated dilution as the minimum removal rate, the maximum effective production rate that can sustain this concentration is *P_syn,max_*(*λ*) *λ*. Converting this effective rate to the corresponding designed production rate using the synthesis multiplier defined above yields the growth-dependent ceiling on the designed production rate:

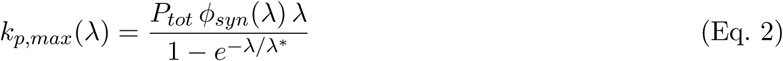

where

*k_p,max_*(*λ*): Maximum feasible designed production rate at growth rate *λ* (*µM/h*) *P_tot_*: Total cellular protein concentration in exponentially growing *E. coli* (*mM*)
*ϕ_syn_*(*λ*): Maximum proteome fraction available for synthetic protein expression at growth rate *λ*
*λ*: Specific growth rate of the host (*h^−^*^1^)
*λ^∗^*: Growth rate constant of the synthesis multiplier (*h^−^*^1^)

The feasible production rate space at a given growth rate was bounded by the line *k_p_*_1_ + *k_p_*_2_ = *k_p,max_*(*λ*), and the robustness metric was computed as the bi-stable fraction of this feasible region. The values of the proteome-partitioning parameters (*ϕ_hk_*, *ϕ_r_*_0_, *w_r_*, *P_tot_*) and their derivation from previously reported measurements are provided in Supplementary Methods and Table S1.

### 5.3 Dynamical modeling framework

In order to systematically evaluate the effects of perturbing the model parameters on the switch’s function, we formulated a general dynamical modeling and paired it with a previously described continuity-analysis framework that accepts as its inputs an ordinary differential equation model of any gene regulatory motif - or more generally, any dynamical process - together with analytical expressions for its Jacobian and nullclines. Given a set of parameter values, the framework computes all the stable and unstable steady states of the model, the separatrices dividing the basins of attraction in multi-stable systems, and, where multiple stable states exist, the time required to switch between them upon the addition of an effector molecule (e.g. IPTG or aTc in the case of the genetic toggle switch). The framework is packaged as a Python module, dynaCA, which uses functionalities from the scipy^39^ package for root finding and numerical integration and couples to the PyDSTool ^40^ package for continuation analysis. The module is available on GitHub ^41^.

#### Steady states and their stability

For a given parameter set, the steady states were located by solving the system *f* (**x**; *θ*) = **0**, where *f* is the vector of model equations, **x** the vector of repressor concentrations, and *θ* the model parameters. Because the system is multi-stable, a single initial guess is insufficient to recover all steady states; we therefore performed multi-start root finding, seeding the solver with a mesh of initial points spanning each state variable geometrically from zero up to an order of magnitude above its maximal attainable concentration (*k_p_/k_deg_*). Each seed was solved using the hybrid Powell method (`scipy.optimize.root`) with the analytical Jacobian supplied, and the resulting roots were de-duplicated to yield the distinct steady states. The stability of each steady state was determined from the eigenvalues of the Jacobian evaluated at that point: a state was classified as stable if all eigenvalues had negative real parts, and unstable otherwise. A bi-stable configuration corresponds to two stable steady states separated by a single unstable steady state.

#### Robustness metric via two-parameter continuation

The robustness metric was computed by delimiting the region of the repressor production-rate space (*k_p_*_1_, *k_p_*_2_) over which the motif is bi-stable, using continuation analysis in `PyDSTool`. For a fixed set of the remaining parameters, the framework first located the equilibria of the system and, by tracking the eigenvalues of the Jacobian along a one-parameter sweep of a production rate, identified the limit points - the fold bifurcations at which a pair of steady states is created or annihilated, and hence the points at which the motif transitions between bi-stable and mono-stable behavior. Starting from one such limit point, two-parameter continuation was then performed in *k_p_*_1_ and *k_p_*_2_ simultaneously to trace the entire locus of limit points bounding the bi-stable region in the production-rate plane. The robustness metric was computed as the fraction of the feasible production-rate space enclosed by this locus - i.e. the fraction of (*k_p_*_1_*, k_p_*_2_) combinations yielding two stable and one unstable steady state - where the feasible space is bounded by the growth-dependent production ceiling *k_p_*_1_ + *k_p_*_2_ *≤ k_p,max_*(*λ*) (Methods 5.2).

#### Speed metric

The speed of switching was determined by simulating the transition between stable states following the addition of inducer. Inducer addition was modeled by setting the dissociation constant of the corresponding repressor to a very large value (*K_diss_ → ∞*), reflecting sequestration of the repressor by the inducer and its consequent inability to bind its promoter target. Throughout this work we consider the fast-switching regime, in which the inducer is assumed to be present continuously until the new steady state is reached. Starting from the initial stable state, the model equations were integrated forward in time using an implicit Radau scheme (`scipy.integrate.solve_ivp`), and the switch time was defined as the time at which both repressor concentrations had reached their new steady-state values, taken as the point at which the normalized deviation of each concentration from its final value fell below a fixed threshold. Because the switch time can vary with the repressor production rates within the bi-stable region, whereas the robustness metric integrates over all production rates, we evaluated the switch time at the four production-rate extrema of the limit-point locus obtained from the robustness analysis and defined the speed metric as the inverse of the average of these four switch times.

#### Nullclines and separatrices

For visualization of the phase plane (Figure S2), the nullclines were evaluated directly from the supplied nullcline expressions over a grid of concentrations, and the separatrix dividing the two basins of attraction was computed by integrating the dynamics in negative time from the unstable steady state - starting from small perturbations along its stable eigenvectors in both directions - so that the backward trajectories converge onto the separatrix. The nullclines and separatrices were used only to illustrate the structure of the phase plane and did not enter the computation of the robustness or speed metrics.

### 5.4 Design and construction of DNA parts and plasmids

The RBS calculator (v2.1) proposed by Reis et al ^22^ was used to design constructs with varying translation initiation rates (TIRs) or strengths of the Ribosome Binding Sequence (RBS) and to calculate the RBS strengths of previously available constructs, where necessary. The genetic toggle switch constructed by Litcofsky et al ^13^ was used as the template to construct the fusion toggle switch, which was then used as the base plasmid for all further constructions. Unique Nucleotide Sequences (UNSs) ^42^ were added upstream and downstream of each genetic part to improve their modularity and enable rapid assembly of different combinations of genetic parts. Preliminary DNA parts for varying RBS’s and protease affinity tags of the repressors were synthesized by Ranomics Inc. Synthetic oligonucleotide parts for DNA assembly reactions were synthesized by Eurofins Scientific and Integrated DNA Technologies Inc. Ligase Cycling Reaction (LCR)^37^ and Gibson assembly ^43^ were used to assemble different DNA parts into complete plasmids. Plasmids were maintained in *E. coli* (DH10β) cells after assembly by transfection through heat shock or electroporation. An Opentrons robotic liquid handler was programmed to perform DNA and plasmid extractions where higher throughput was required. All PCR amplifications for plasmid construction were performed using Q5 High-Fidelity Polymerase (NEB). All plasmids were sequence verified through Sanger sequencing. *E. coli* cells harboring the target plasmids were stored in 20% glycerol at -80°C. Sequence files for plasmids used in this study can be found attached as supplementary files and are described in Table S6.

### 5.5 Characterization of fusion protein constructs and toggle switch variants

The constructed plasmids were transformed into *E. coli* (MG1655) Δ lacI - the target host for final characterization, through electroporation. All strains were characterized using an in-house microplate based high throughput robotic platform that has been previously described ^38^. Cells were grown in microtiter plates overnight using LB media supplemented with 1% glucose and the appropriate inducer (2 mM isopropyl β-D-1-thiogalactopyranoside - IPTG or 500 ng/mL anhydrotetracycline - aTc). Following this, they were transferred to fresh Rich Defined Media with the same inducer, supplemented with 1% glucose and allowed to adapt to the new media overnight. Rich defined media (RDM) was used as the growth medium for phenotyping experiments. It is composed of a carbon source (D-glucose at various concentrations), salts (3.5 g/L KH_2_PO_4_, 5 g/L K_2_HPO_4_, 3.5 g/L (NH_4_)_2_HPO_4_, 1 mM MgSO_4_, 0.1mM CaCl_2_), 1 mM 3-morpholinopropane-1-sulfonic acid (MOPS), amino acid supplements (0.8 mM alanine, 5.2 mM arginine, 0.4 mM aspargine, 0.4 mM aspartate, 0.1 mM cysteine, 0.6 mM glutamate, 0.6 mM glutamine, 0.8 mM glycine, 0.2 mM histidine, 0.4 mM isoleucine, 0.8 mM leucine, 0.4 mM lysine, 0.2 mM methionine, 0.4 mM phenylalanine, 0.4 mM proline, 10 mM serine, 0.4 mM threonine, 0.1 mM tryptophan, 0.2 mM tyrosine, and 0.6 mM valine), nucleotide supplements (0.1 mM each of adenine, cytosine, guanine, and uracil), and vitamin supplements (0.01 mM each of thiamine, calcium pantothenate, *p*-aminobenzoic acid, *p*-hydroxybenzoic acid, and 2,3-dihydroxybenzoic acid) - adapted from the defined media composition described previously ^44^.

To examine the speed of switching, overnight RDM cultures were then washed with RDM media without any inducer to remove traces of the previously used inducer and transferred to fresh rich defined media ^44^ at a uniform cell density corresponding to *OD*_700_ of 0.01. The required inducer (2 mM IPTG to strains grown overnight on aTc and 500 ng/mL aTC to strains grown overnight on IPTG) was added to the plates which were then incubated in a Tecan Spark microplate reader at 37*^◦^*C. The cell densities and concentrations of reporter proteins were monitored by recording absorbance at 700 nm and fluorescence intensities using the following parameters: GFP (Excitation: 488 nm, Emission: 525 nm, Gain: 90), mCherry (Excitation: 570 nm, Emission: 610 nm, Gain: 110). For bi-stability tests, overnight cultures of *E. coli* in RDM containing the appropriate inducer were washed with media lacking inducer to remove all traces of the previously used inducer. Then, cells were inoculated in fresh RDM without inducer at a cell density of OD_700_ = 0.01 and grown to saturation. Following this, the cells were progressively transferred two more times to fresh media lacking inducer at a 1:100 dilution at 12 hour intervals, with the cell density and fluorescent reporter concentrations being recorded at the end of each transfer.

### 5.6 Data analysis

Blank subtracted fluorescence values measured in a microplate reader, normalized to the measured cell density values were used to track repressor protein levels in the cells. For GFP measurements, the auto-fluorescence of *E. coli* was further subtracted prior to normalization to the cell densities. The auto-fluorescence of *E. coli* lacking GFP was measured at different OD values for this purpose. For the switch speed experiments, the time required to switch was calculated as the time taken for the fluorescent reporters to reach their steady state concentrations. This was done by approximating the first derivative of OD normalized fluorescence data and determining the time at which its absolute value becomes lower than a fixed threshold. For bi-stability analysis, the GFP and mCherry fluorescence at the end of each transfer was recorded. The cells were considered stable in their state if the fluorescence values of the corresponding reporter was at a detectable level after each of the three transfers, indicating that the corresponding repressor is also still at high levels. A replicate was classified as bi-stable if the reporter corresponding to each induced state remained detectable after all three inducer-free passages, mono-stable towards a given state if only that state’s reporter remained detectable, and unstable if neither reporter remained detectable. Each strain was then classified based on its replicates: bi-stable if all replicates were bi-stable, mono-stable towards a state if any replicate was mono-stable towards that state, and unstable if replicates were mono-stable towards different states or if any replicate was unstable. Data analyses for all sections were conducted using Python on Jupyter notebooks. The jupyter notebooks used to generate figures and process data in this work have been made available on GitHub ^45^. The python based data analysis library - pandas and plotting library - plotly were used extensively for all data analysis and visualization pipelines in this work ^46,47^. Microbial phenotypic data and growth curves were analyzed using the IMPACT framework ^48^.

## Acknowledgements

This work was supported by the Natural Sciences and Engineering Research Council (NSERC), the NSERC Industrial Biocatalysis Network, the Ontario Ministry of Research and Innovation, and Genome Canada.

We thank Dr. James Collins (Massachusetts Institute of Technology, USA) for providing the toggle switch backbone plasmid - pKDL071 and the host strain MG1655 ΔlacI.

## References

[1] Timo Hardiman, Karin Lemuth, Markus A. Keller, Matthias Reuss, and Martin Siemann-Herzberg. Topology of the global regulatory network of carbon limitation in Escherichia coli. J. Biotechnol., 132(4):359–374, 2007. ISSN 01681656. doi: 10.1016/j.jbiotec.2007.08.029. URL https://www.sciencedirect.com/science/article/pii/S0168165607015210.

[2] Joe Lutkenhaus. The regulation of bacterial cell division: A time and place for it. Curr. Opin. Microbiol., 1(2):210–215, 1998. ISSN 13695274. doi: 10.1016/S1369-5274(98)80013-1. URL https://www.sciencedirect.com/science/article/pii/S1369527498800131.

[3] Andrea Pitzschke, Adam Schikora, and Heribert Hirt. MAPK cascade signalling networks in plant defence. Curr. Opin. Plant Biol., 12(4):421–426, 2009. ISSN 13695266. doi: 10.1016/j.pbi.2009.06.008. URL https://www.sciencedirect.com/science/article/pii/S1369526609000715.

[4] Uri Alon. Network motifs: Theory and experimental approaches. Nat. Rev. Genet., 8(6):450–461, 2007. ISSN 14710056. doi: 10.1038/nrg2102.

[5] James E. Ferrell. Feedback regulation of opposing enzymes generates robust, all-or-none bistable responses. Curr. Biol., 18(6):R244–R245, 2008. ISSN 09609822. doi: 10.1016/j.cub.2008.02.035. URL https://www.sciencedirect.com/science/article/pii/S0960982208001814.

[6] Timothy S. Gardner, Charles R. Cantor, and James J. Collins. Construction of a genetic toggle switch in Escherichia coli. Nature, 403(6767):339–342, 2000. ISSN 00280836. doi: 10.1038/35002131. URL http://www.nature.com/doifinder/10.1038/35002131.

[7] Naveen Venayak, Kaushik Raj, Rohil Jaydeep, and Radhakrishnan Mahadevan. An Optimized Bistable Metabolic Switch to Decouple Phenotypic States during Anaerobic Fermentation. ACS Synth. Biol., 7(12):2854–2866, oct 2018. ISSN 21615063. doi: 10.1021/acssynbio.8b00284.

[8] William Bothfeld, Grace Kapov, and Keith E.J. Tyo. A Glucose-Sensing Toggle Switch for Autonomous, High Productivity Genetic Control. ACS Synth. Biol., 6(7):1296–1304, jul 2017. ISSN 21615063. doi: 10.1021/acssynbio.6b00257. URL http://pubs.acs.org/doi/pdf/10.1021/acssynbio.6b00257 http://pubs.acs.org/doi/abs/10.1021/acssynbio.6b00257.

[9] David T. Riglar, Tobias W. Giessen, Michael Baym, S. Jordan Kerns, Matthew J. Niederhuber, Roderick T. Bronson, Jonathan W. Kotula, Georg K. Gerber, Jeffrey C. Way, and Pamela A. Silver. Engineered bacteria can function in the mammalian gut long-term as live diagnostics of inflammation. Nat. Biotechnol., 35(7):653–658, may 2017. ISSN 15461696. doi: 10.1038/nbt.3879. URL https://www.nature.com/articles/nbt.3879.

[10] Jonathan W. Kotula, S. Jordan Kerns, Lev A. Shaket, Layla Siraj, James J. Collins, Jeffrey C. Way, and Pamela A. Silver. Programmable bacteria detect and record an environmental signal in the mammalian gut. Proc. Natl. Acad. Sci. U. S. A., 111(13):4838–4843, 2014. ISSN 10916490. doi: 10.1073/pnas.1321321111. URL http://www.pnas.org.gate1.inist.fr/content/111/13/4838.long.

[11] Clement T.Y. Chan, Jeong Wook Lee, D. Ewen Cameron, Caleb J. Bashor, and James J. Collins. ’Deadman’ and ’Passcode’ microbial kill switches for bacterial containment. Nat. Chem. Biol., 12(2): 82–86, feb 2016. ISSN 15524469. doi: 10.1038/nchembio.1979. URL http://www.ncbi.nlm.nih.gov/pubmed/26641934 http://www.pubmedcentral.nih.gov/articlerender.fcgi?artid=PMC4718764.

[12] Finn Stirling, Lisa Bitzan, Samuel O’Keefe, Elizabeth Redfield, John W.K. Oliver, Jeffrey Way, and Pamela A. Silver. Rational Design of Evolutionarily Stable Microbial Kill Switches. Mol. Cell, 68(4):686–697.e3, nov 2017. ISSN 10974164. doi: 10.1016/j.molcel.2017.10.033. URL http://www.ncbi.nlm.nih.gov/pubmed/29149596 http://www.pubmedcentral.nih.gov/articlerender.fcgi?artid=PMC5812007.

[13] Kevin D. Litcofsky, Raffi B. Afeyan, Russell J. Krom, Ahmad S. Khalil, and James J. Collins. Iterative plug-and-play methodology for constructing and modifying synthetic gene networks. Nat. Methods, 9(11):1077–1080, nov 2012. ISSN 15487091. doi: 10.1038/nmeth.2205. URL http://www.pubmedcentral.nih.gov/articlerender.fcgi?artid=3492501&tool=pmcentrez&rendertype=abstract.

[14] Jeong Wook Lee, Andras Gyorgy, D. Ewen Cameron, Nora Pyenson, Kyeong Rok Choi, Jeffrey C. Way, Pamela A. Silver, Domitilla Del Vecchio, and James J. Collins. Creating Single-Copy Genetic Circuits. Mol. Cell, 63(2):329–336, jul 2016. ISSN 10974164. doi: 10.1016/j.molcel.2016.06.006. URL http://www.ncbi.nlm.nih.gov/pubmed/27425413 http://www.pubmedcentral.nih.gov/articlerender.fcgi?artid=PMC5407370.

[15] Kaushik Raj, Naveen Venayak, and Radhakrishnan Mahadevan. Novel two-stage processes for optimal chemical production in microbes. Metab. Eng., 62:186–197, nov 2020. ISSN 10967184. doi: 10.1016/j.ymben.2020.08.006.

[16] Jordan J. Sickle, Congjian Ni, Daniel Shen, Zewei Wang, Matthew Jin, and Ting Lu. Integrative Circuit-Host Modeling of a Genetic Switch in Varying Environments. Sci. Rep., 10(1):1–9, may 2020. ISSN 20452322. doi: 10.1038/s41598-020-64921-5. URL https://www.nature.com/articles/s41598-020-64921-5.

[17] Rong Zhang, Jiao Li, Juan Melendez-Alvarez, Xingwen Chen, Patrick Sochor, Hanah Goetz, Qi Zhang, Tian Ding, Xiao Wang, and Xiao Jun Tian. Topology-dependent interference of synthetic gene circuit function by growth feedback. Nat. Chem. Biol., 16(6):695–701, apr 2020. ISSN 15524469. doi: 10.1038/s41589-020-0509-x. URL https://www.nature.com/articles/s41589-020-0509-x.

[18] Stefan Klumpp, Zhongge Zhang, and Terence Hwa. Growth rate-dependent global effects on gene expression in bacteria. Cell, 139(7):1366–75, dec 2009. ISSN 1097-4172. doi: 10.1016/j.cell.2009.12.001. URL http://www.ncbi.nlm.nih.gov/pubmed/20064380 http://www.pubmedcentral.nih.gov/articlerender.fcgi?artid=PMC2818994.

[19] M Brenowitz, N Mandal, A Pickar, E Jamison, and S Adhya. Dna-binding properties of a lac repressor mutant incapable of forming tetramers. Journal of Biological Chemistry, 266(2):1281–1288, Jan 1991. doi: 10.1016/s0021-9258(17)35313-9.

[20] Robert Daber, Matthew A. Sochor, and Mitchell Lewis. Thermodynamic analysis of mutant lac repressors. Journal of Molecular Biology, 409(1):76–87, May 2011. doi: 10.1016/j.jmb.2011.03.057.

[21] Nicholas Benson, Curtis Adams, and Philip Youderian. Genetic selection for mutations that impair the co-operative binding of lambda repressor. Molecular Microbiology, 11(3):567–579, Feb 1994. doi: 10.1111/j.1365-2958.1994.tb00337.x.

[22] Alexander C. Reis and Howard M. Salis. An automated model test system for systematic development and improvement of gene expression models. ACS Synth. Biol., 9(11):3145–3156, 2020. ISSN 21615063. doi: 10.1021/acssynbio.0c00394. URL https://pubs.acs.org/doi/full/10.1021/acssynbio.0c00394.

[23] Kathleen E. McGinness, Tania A. Baker, and Robert T. Sauer. Engineering Controllable Protein Degradation. Mol. Cell, 22(5):701–707, 2006. ISSN 10972765. doi: 10.1016/j.molcel.2006.04.027.

[24] Rolf Lutz and Hermann Bujard. Independent and tight regulation of transcriptional units in escherichia coli via the lacr/o, the tetr/o and arac/i1-i2 regulatory elements. Nucleic Acids Research, 25(6):1203– 1210, 03 1997. ISSN 0305-1048. doi: 10.1093/nar/25.6.1203. URL https://doi.org/10.1093/nar/25.6.1203.

[25] Michael B. Elowitz and Stanislas Leibier. A synthetic oscillatory network of transcriptional regulators. Nature, 403(6767):335–338, 2000. ISSN 00280836. doi: 10.1038/35002125.

[26] David W. Erickson, Severin J. Schink, Vadim Patsalo, James R. Williamson, Ulrich Gerland, and Terence Hwa. A global resource allocation strategy governs growth transition kinetics of Escherichia coli. Nature, 551(7678):119, nov 2017. ISSN 14764687. doi: 10.1038/NATURE24299. URL /pmc/articles/PMC5901684//pmc/articles/PMC5901684/?report=abstract https://www.ncbi.nlm.nih.gov/pmc/articles/PMC5901684/.

[27] Marius Hintsche and Stefan Klumpp. Dilution and the theoretical description of growth-rate dependent gene expression. Journal of Biological Engineering, 7(1):22, 2013. doi: 10.1186/1754-1611-7-22.

[28] Matthew Scott, Carl W. Gunderson, Eduard M. Mateescu, Zhongge Zhang, and Terence Hwa. Interdependence of cell growth and gene expression: Origins and consequences. Science, 330(6007): 1099–1102, Nov 2010. doi: 10.1126/science.1192588.

[29] Matteo Mori, Terence Hwa, Olivier C. Martin, Andrea De Martino, and Enzo Marinari. Constrained allocation flux balance analysis. PLOS Computational Biology, 12(6), Jun 2016. doi: 10.1371/journal.pcbi.1004913.

[30] Jia Xu and Kathleen S. Matthews. Flexibility in the inducer binding region is crucial for allostery in the Escherichia coli lactose repressor. Biochemistry, 48(22):4988–4998, jun 2009. ISSN 00062960. doi: 10.1021/bi9002343. URL https://pubs.acs.org/doi/full/10.1021/bi9002343.

[31] Subhayu Basu, Yoram Gerchman, Cynthia H. Collins, Frances H. Arnold, and Ron Weiss. A synthetic multicellular system for programmed pattern formation. Nature, 434(7037):1130–1134, apr 2005. ISSN 00280836. doi: 10.1038/nature03461. URL http://www.ncbi.nlm.nih.gov/pubmed/15858574.

[32] Jiunn R. Chen. Improving the speed of the genetic toggle switch without sacrificing its dynamic stability. *Phys. Rev. E - Stat. Nonlinear*, Soft Matter Phys., 73(4):041901, apr 2006. ISSN 15502376. doi: 10.1103/PhysRevE.73.041901. URL http://link.aps.org/doi/10.1103/PhysRevE.73.041901.

[33] Daniel Huang, William J. Holtz, and Michel M. Maharbiz. A genetic bistable switch utilizing nonlinear protein degradation. J. Biol. Eng., 6(1):9, 2012. ISSN 17541611. doi: 10.1186/1754-1611-6-9.

34. F.C. Neidhardt. Escherichia Coli and Salmonella: Cellular and Molecular Biology. Number v. 1-2 in Escherichia Coli and Salmonella: Cellular and Molecular Biology. ASM Press, 1996. ISBN 9781555810849. URL https://books.google.ca/books?id=SPSxzgEACAAJ.

[35] Xiaoying Chen, Jennica L. Zaro, and Wei Chiang Shen. Fusion protein linkers: Property, design and functionality. Adv. Drug Deliv. Rev., 65(10):1357–1369, 2013. ISSN 0169409X. doi: 10.1016/j.addr.2012.09.039. URL https://www.sciencedirect.com/science/article/pii/S0169409X12003006.

[36] Jianshen Hou, Cong Gao, Liang Guo, Jens Nielsen, Qiang Ding, Wenxiu Tang, Guipeng Hu, Xiulai Chen, and Liming Liu. Rewiring carbon flux in Escherichia coli using a bifunctional molecular switch. Metab. Eng., 61:47–57, 2020. ISSN 10967184. doi: 10.1016/j.ymben.2020.05.004. URL https://www.sciencedirect.com/science/article/pii/S1096717620300860.

[37] Stefan De Kok, Leslie H. Stanton, Todd Slaby, Maxime Durot, Victor F. Holmes, Kedar G. Patel, Darren Platt, Elaine B. Shapland, Zach Serber, Jed Dean, Jack D. Newman, and Sunil S. Chandran. Rapid and reliable DNA assembly via ligase cycling reaction. ACS Synth. Biol., 3(2):97–106, 2014. ISSN 21615063. doi: 10.1021/sb4001992.

[38] Kaushik Raj, Naveen Venayak, Patrick Diep, Sai Akhil Golla, Alexander F. Yakunin, and Radhakrishnan Mahadevan. Automation assisted anaerobic phenotyping for metabolic engineering. Microb. Cell Fact., 20(1):1–16, 2021. ISSN 14752859. doi: 10.1186/s12934-021-01675-3. URL https://microbialcellfactories.biomedcentral.com/articles/10.1186/s12934-021-01675-3.

[39] Pauli Virtanen, Ralf Gommers, Travis E. Oliphant, Matt Haberland, Tyler Reddy, David Cournapeau, Evgeni Burovski, Pearu Peterson, Warren Weckesser, Jonathan Bright, Stéfan J. van der Walt, Matthew Brett, Joshua Wilson, K. Jarrod Millman, Nikolay Mayorov, Andrew R.J. Nelson, Eric Jones, Robert Kern, Eric Larson, C. J. Carey, İlhan Polat, Yu Feng, Eric W. Moore, Jake VanderPlas, Denis Laxalde, Josef Perktold, Robert Cimrman, Ian Henriksen, E. A. Quintero, Charles R. Harris, Anne M. Archibald, Antônio H. Ribeiro, Fabian Pedregosa, Paul van Mulbregt, Aditya Vijaykumar, Alessandro Pietro Bardelli, Alex Rothberg, Andreas Hilboll, Andreas Kloeckner, Anthony Scopatz, Antony Lee, Ariel Rokem, C. Nathan Woods, Chad Fulton, Charles Masson, Christian Häggström, Clark Fitzgerald, David A. Nicholson, David R. Hagen, Dmitrii V. Pasechnik, Emanuele Olivetti, Eric Martin, Eric Wieser, Fabrice Silva, Felix Lenders, Florian Wilhelm, G. Young, Gavin A. Price, Gert Ludwig Ingold, Gregory E. Allen, Gregory R. Lee, Hervé Audren, Irvin Probst, Jörg P. Dietrich, Jacob Silterra, James T. Webber, Janko Slavič, Joel Nothman, Johannes Buchner, Johannes Kulick, Johannes L. Schönberger, José Vinícius de Miranda Cardoso, Joscha Reimer, Joseph Harrington, Juan Luis Cano Rodríguez, Juan Nunez-Iglesias, Justin Kuczynski, Kevin Tritz, Martin Thoma, Matthew Newville, Matthias Kümmerer, Maximilian Bolingbroke, Michael Tartre, Mikhail Pak, Nathaniel J. Smith, Nikolai Nowaczyk, Nikolay Shebanov, Oleksandr Pavlyk, Per A. Brodtkorb, Perry Lee, Robert T. McGibbon, Roman Feldbauer, Sam Lewis, Sam Tygier, Scott Sievert, Sebastiano Vigna, Stefan Peterson, Surhud More, Tadeusz Pudlik, Takuya Oshima, Thomas J. Pingel, Thomas P. Robitaille, Thomas Spura, Thouis R. Jones, Tim Cera, Tim Leslie, Tiziano Zito, Tom Krauss, Utkarsh Upadhyay, Yaroslav O. Halchenko, and Yoshiki Vázquez-Baeza. SciPy 1.0: fundamental algorithms for scientific computing in Python. Nat. Methods, 17(3):261–272, 2020. ISSN 15487105. doi: 10.1038/s41592-019-0686-2.

[40] Robert Clewley. Hybrid models and biological model reduction with pydstool. PLOS Computational Biology, 8(8):1–8, 08 2012. doi: 10.1371/journal.pcbi.1002628. URL https://doi.org/10.1371/journal.pcbi.1002628.

41. Kaushik Raj, William T Z Wong, Beini Zhang, and Radhakrishnan Mahadevan. Novel dynamical modeling framework GitHub repository. Available from: https://github.com/LMSE/dynaCA. Accessed 10 July 2022, 2022.

[42] Joseph P. Torella, Florian Lienert, Christian R. Boehm, Jan Hung Chen, Jeffrey C. Way, and Pamela A. Silver. Unique nucleotide sequence-guided assembly of repetitive DNA parts for synthetic biology applications. Nat. Protoc., 9(9):2075–2089, 2014. ISSN 17502799. doi: 10.1038/nprot.2014.145.

[43] Daniel G. Gibson, Lei Young, Ray Yuan Chuang, J. Craig Venter, Clyde A. Hutchison, and Hamilton O. Smith. Enzymatic assembly of DNA molecules up to several hundred kilobases. Nat. Methods, 6(5):343–345, may 2009. ISSN 15487091. doi: 10.1038/nmeth.1318. URL http://www.nature.com/doifinder/10.1038/nmeth.1318.

[44] F. C. Neidhardt, P. L. Bloch, and D. F. Smith. Culture medium for enterobacteria. J. Bacteriol., 119 (3):736–747, 1974. ISSN 00219193. doi: 10.1128/jb.119.3.736-747.1974. URL https://pubmed.ncbi.nlm.nih.gov/4604283/.

45. Kaushik Raj, William T Z Wong, Beini Zhang, and Radhakrishnan Mahadevan. Enhanced bi-stable motifs GitHub repository. Available from: https://github.com/LMSE/enhanced_bistable_ switches. Accessed 10 July 2022, 2022.

46. Pandas Development team. pandas-dev/pandas: Pandas, feb 2020.

47. Plotly Technologies Inc. Collaborative data science, 2015. URL https://plot.ly.

[48] Naveen Venayak, Kaushik Raj, and Radhakrishnan Mahadevan. Impact framework: A python package for writing data analysis workflows to interpret microbial physiology. Metab. Eng. Commun., 9, 2019. ISSN 22140301. doi: 10.1016/j.mec.2019.e00089.

