## Supplemental Information for "Design principles for bistable genetic switches emerge from a growth-dependent speed–robustness trade-off"

### S1 Supplementary Methods

#### S1.1 Determination of theoretical productivity loss due to slow switching

To quantify the impact of switch delay on bioprocess productivity, we simulated a two-stage chemical production process in *E. coli* comprising a growth and a production stage using dynamic flux balance analysis (dFBA) with the iML1515 genome-scale metabolic model of *E. coli*[1]. The procedure used here follows that described previously[2].

We considered two models of substrate uptake. In the first, the substrate uptake rate was held fixed at 10 mmol/gDW.h. In the second, the substrate uptake rate followed a logistic function with increasing growth rate, as described previously[2]. For each uptake assumption, a production envelope was constructed for five representative products spanning a range of carbon numbers - acetate, ethanol, D-lactate, succinate, and citrate by determining the achievable product flux at each value of the growth rate between zero and the maximum growth rate.

The bioprocess was initiated with 0.1 g/L biomass and 500 mmol/L of glucose as the substrate. The process was modelled as a transition from a growth stage, in which cellular resources are directed towards biomass formation, to a production stage, in which they are directed towards synthesis of the target product. The switch delay was defined as the time taken to transition between these two stages. For a given switch delay, the optimal switch time and the corresponding combination of substrate, product, and growth fluxes in each stage were determined so as to maximize the bioprocess productivity, defined as the product titer divided by the total fermentation time. This optimization was performed using the `minimize` function of the `scipy.optimize` package[3] with the COBYLA method. The resulting optimal productivity was computed for switch delays ranging from 0 to 25 h and is plotted against the switch delay for each product and each uptake assumption in Figure S1.

Across all five products, the maximum achievable productivity decreased monotonically with increasing switch delay under both substrate uptake assumptions, confirming that the productivity cost of slow switching is preserved regardless of how substrate uptake is modelled.

#### S1.2 Estimation of proteome-partitioning parameters

The growth-dependent ceiling on the designed production rate (Main Text Methods 5.2) requires numerical values for the housekeeping proteome fraction $\phi_{hk}$, the ribosomal proteome fraction parameters $\phi_{r0}$ and $w_r$, and the total cellular protein concentration $P_{tot}$. These were derived from previously reported measurements as described below.

**Ribosomal fraction:** For exponentially growing *E. coli*, the RNA-to-protein ratio $r$ increases linearly with the specific growth rate $\lambda$, following

$$r = r_0 + \frac{\lambda}{k_t} \qquad \text{(Eq. S1)}$$

where $r_0$ is the intercept and $k_t$ is the inverse of the slope[4]. We used the reported values $r_0 = 0.087$ and $k_t = 4.5\, h^{-1}$ as described previously[4]. Since the RNA in these cells is predominantly ribosomal, $r$ serves as a proxy for ribosomal content. To convert the RNA-to-protein ratio into a ribosomal protein fraction, we applied the conversion factor $\rho = 0.76\, \mu g$ protein/$\mu g$ RNA relating ribosomal RNA to ribosomal protein[4]:

$$\phi_r(\lambda) = \rho\, r = \rho \left( r_0 + \frac{\lambda}{k_t} \right) = \phi_{r0} + w_r \lambda \qquad \text{(Eq. S2)}$$

Matching terms, the intercept and slope of the ribosomal protein fraction are $\phi_{r0} = \rho\, r_0$ and $w_r = \rho/k_t$, which evaluate to $\phi_{r0} \approx 0.07$ and $w_r \approx 0.17\, h$ using the values above.

**Housekeeping fraction and total protein:** The housekeeping fraction was taken to be $\phi_{hk} = 0.5$, consistent with proteome-partitioning analyses of *E. coli*[4,5], and the total cellular protein concentration in exponentially growing *E. coli* was taken to be $P_{tot} = 4\,mM$ as reported previously[6].

**Summary of parameter values:** The proteome-partitioning parameter values used to evaluate $k_{p,max}(\lambda)$ are summarized in Table S1.

#### S1.3 Estimation of relative degradation rates

The relative degradation rates of the repressor-reporter fusion constructs (Table S3, Table S4) were estimated from their steady-state fluorescence intensities. Each of the 20 constructs per repressor (5 ribosome binding site strengths $\times$ 4 degradation tags) was grown for 12 hours in the presence of the inducer required to drive its expression, and the endpoint fluorescence intensity was recorded. Fluorescence trajectories were inspected visually to confirm that they had reached steady state by the end of the growth period.

The concentration of a fusion protein $P$ evolves as its production, at rate $k_p$, balanced against its removal by active degradation ($k_{deg}$) and growth-mediated dilution ($\lambda$):

$$\frac{dP}{dt} = k_p - (k_{deg} + \lambda)\,P \qquad \text{(Eq. S3)}$$

assuming repression of its production has been sequestered successfully by addition of an inducer.

At steady state ($dP/dt = 0$), the concentration - and hence the measured fluorescence $F \propto P$ - is given by

$$P_{ss} = \frac{k_p}{k_{deg} + \lambda} \qquad \text{(Eq. S4)}$$

so that steady-state fluorescence is inversely proportional to the effective removal rate ($k_{deg} + \lambda$) at a fixed production rate. Within a given ribosome binding site strength, $k_p$ is fixed, so differences in steady-state fluorescence between degradation tags reflect differences in their effective removal rates: a construct with a faster-degrading tag reaches a lower steady-state fluorescence.

Accordingly, for each ribosome binding site strength we identified the construct with the highest steady-state fluorescence - corresponding to the lowest degradation rate, which in almost all cases was the untagged construct - and used it as the baseline for that strength. In the single case where two constructs had closely comparable fluorescence, the construct with the marginally higher fluorescence (which carried a degradation tag) was taken as the baseline. The estimated relative degradation rate of each construct at a given ribosome binding site strength was then computed as the ratio of the baseline fluorescence to that construct's fluorescence:

$$\left(\frac{k_{deg} + \lambda}{k_{deg,base} + \lambda}\right) = \frac{F_{base}}{F} \qquad \text{(Eq. S5)}$$

which follows directly from Equation Eq. S4 at fixed $k_p$. Because the production rate is held constant only within a given ribosome binding site strength, these estimated relative degradation rates are comparable within each strength but not across different strengths.

#### S2 Supplementary figures

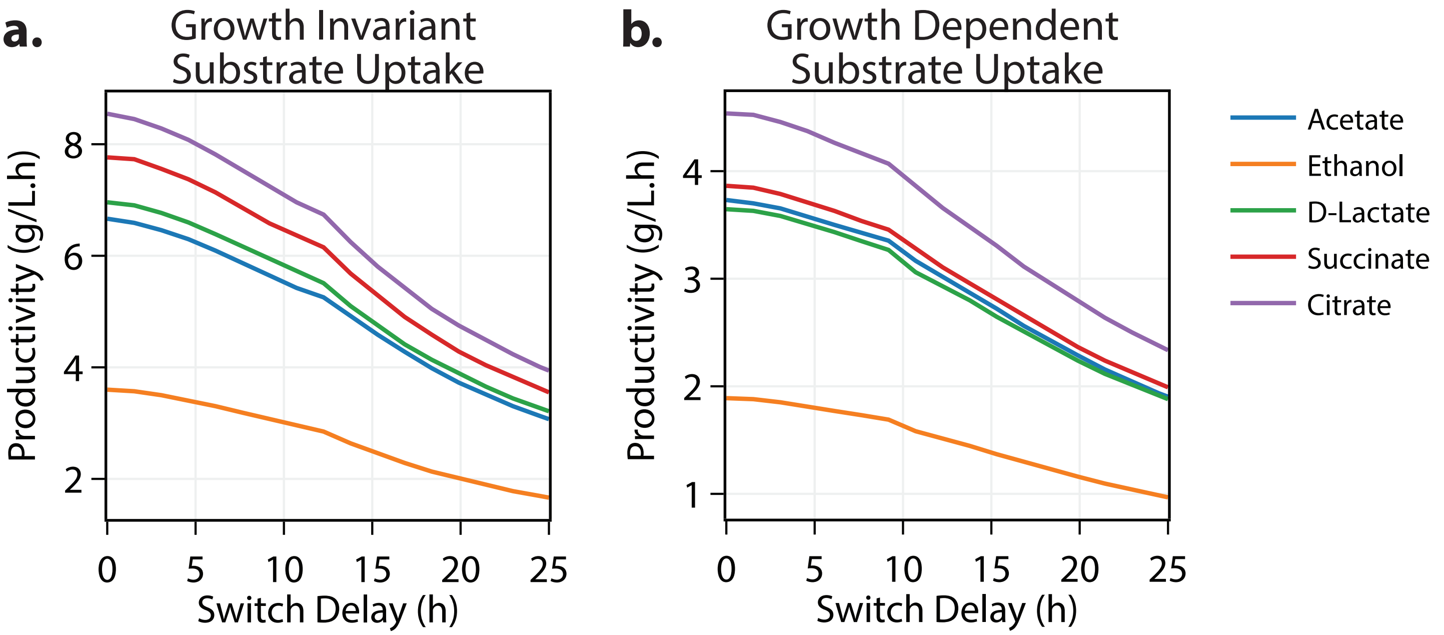


**Figure S1: Productivity declines with switching delay in a simulated two-stage bioprocess.** A dynamic flux balance analysis framework was used to simulate a two-stage chemical production process in *E. coli*, divided into a growth and a production stage using a genome-scale metabolic model. For each value of switch delay (i.e. the time taken to transition from growth to production), the maximum achievable productivity (product titer divided by fermentation time) was determined for five representative products under the assumption that **a.** substrate uptake rate does not change with growth rate, **b.** substrate uptake scales with host growth rates. In both cases, the maximum achievable productivity decreases monotonically with increasing switch delay across all examined products, even though the flux distributions of each stage were free to re-optimize at each delay. Note that the absolute productivities differ between the two uptake assumptions, but the monotonic decline with switch delay is preserved across the assumptions. A full description of the models and methods used for this simulation are available in Methods S1.1.

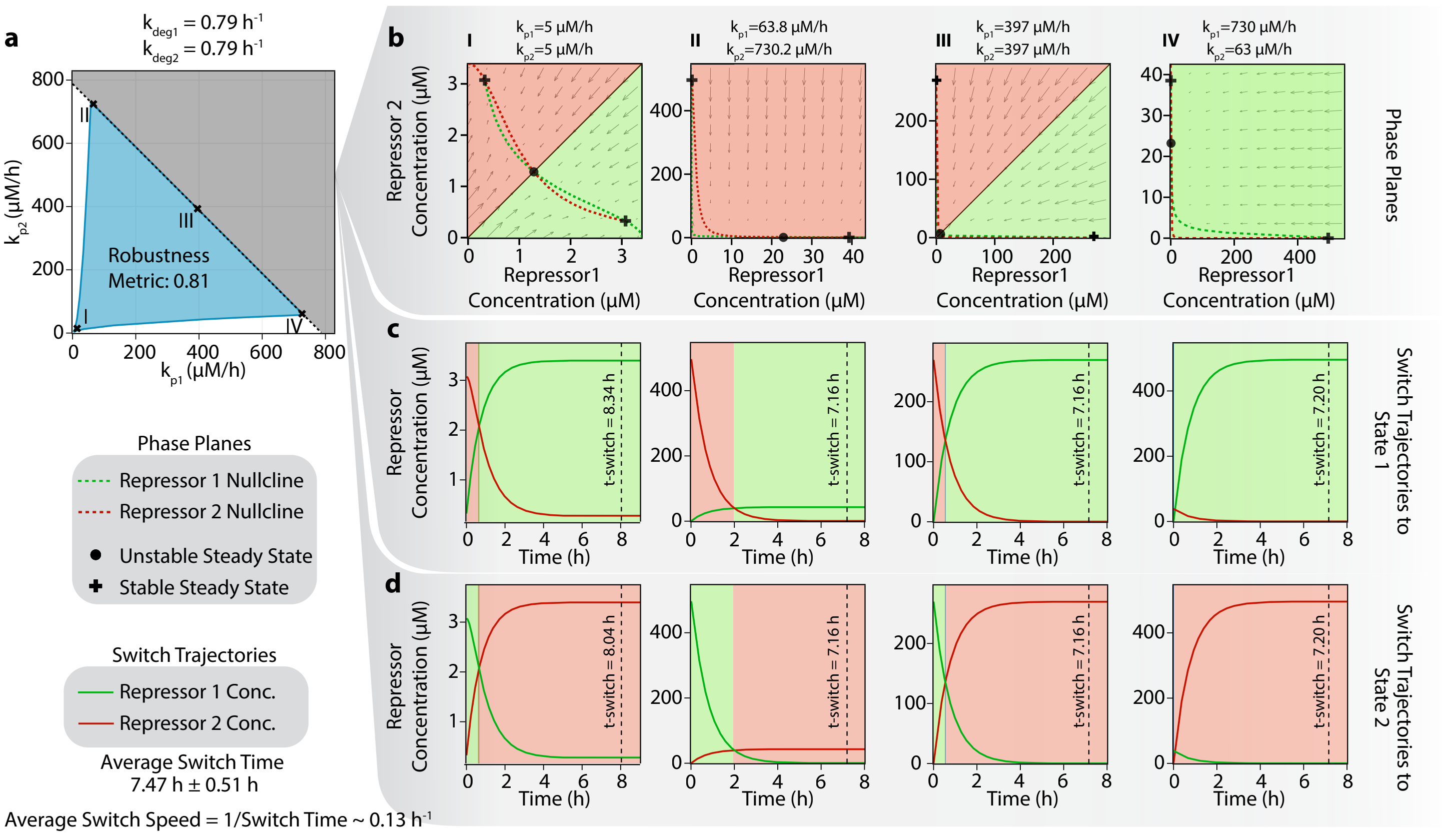


**Figure S2: Determination of Speed Metric: a.** Parameter values from the extrema of the robustness analysis are used to perform a simulation of the ordinary differential equations describing the bi-stable motifs and obtain **b.** the phase plane and **c., d.** the switch trajectories to either state using our dynamical modeling framework. The average time taken to switch to either state is first determined and the switch speed is calculated as the multiplicative inverse of the average switch time.

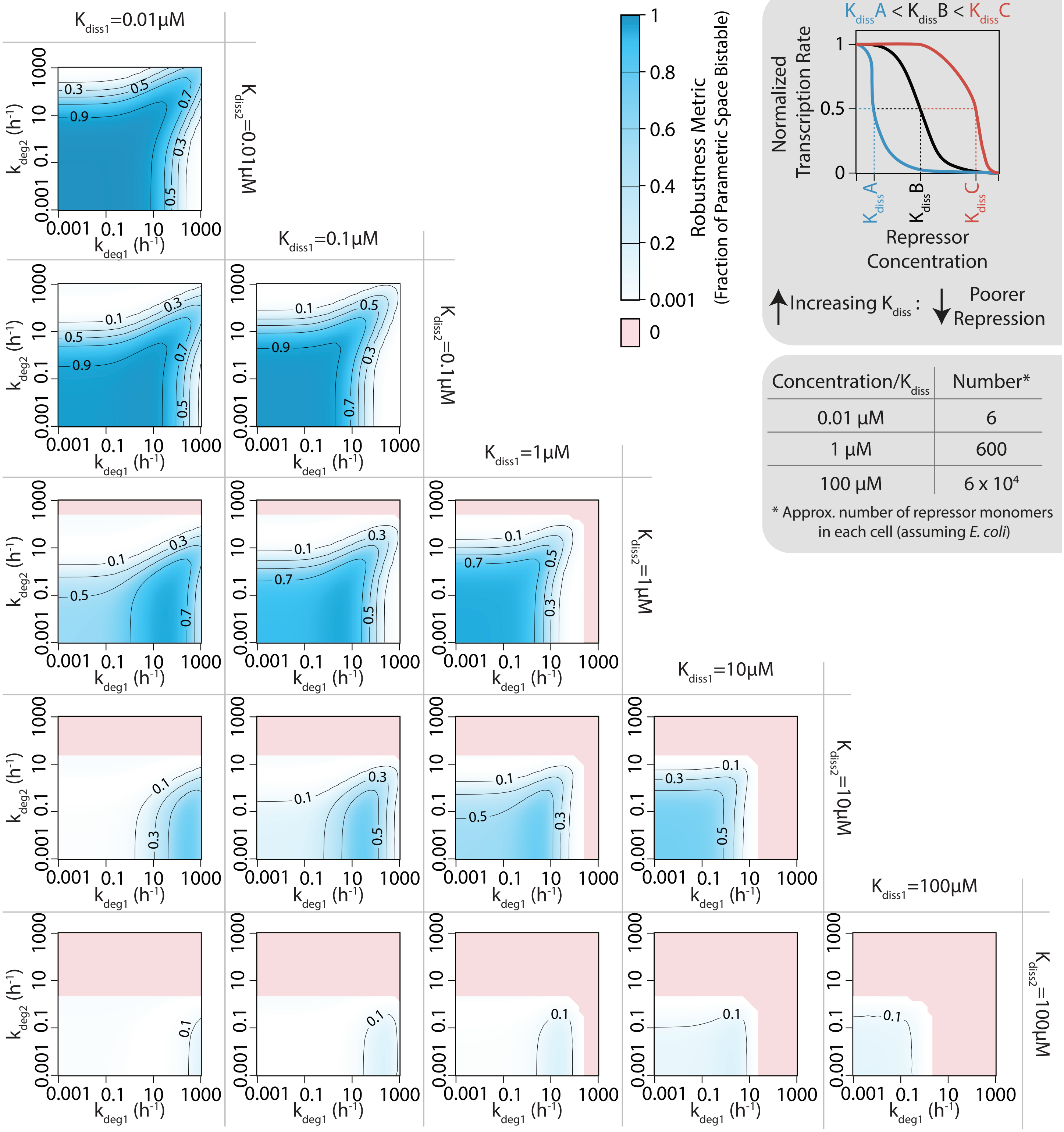


**Figure S3: Robustness landscape for all feasible parameter combinations**. The robustness landscape over all feasible protein degradation rates, calculated for different combinations of $K_{diss}$ values that represent the physical limits of experimentally observable values for the parameter. The shaded boxes to the top-right illustrate the physical significance of the parameter - $K_{diss}$.

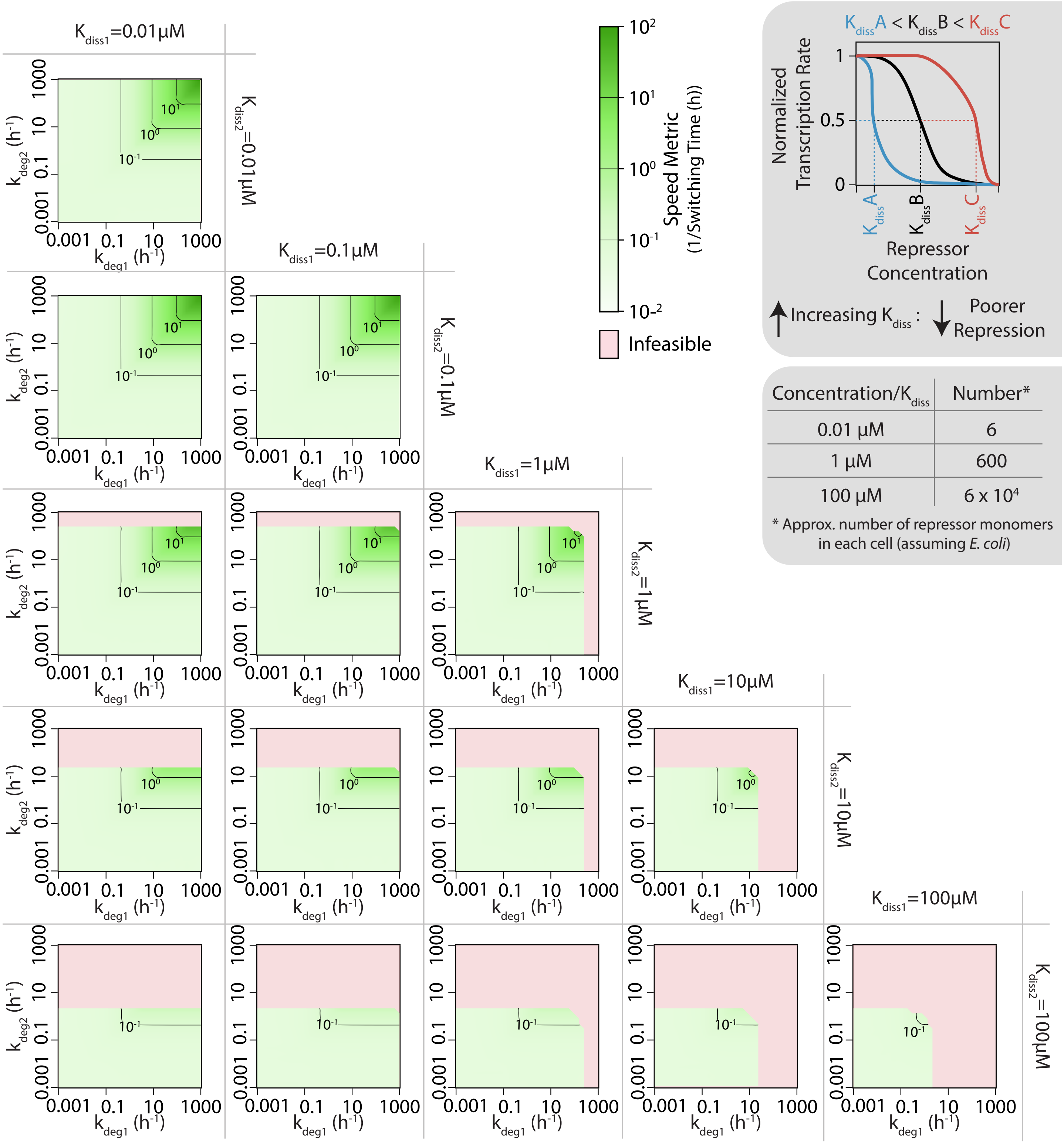


**Figure S4: Speed metric landscape for all feasible parameter combinations**. The robustness landscape over all feasible protein degradation rates, calculated for different combinations of $K_{diss}$ values that represent the physical limits of experimentally observable values for the parameter. The shaded boxes to the top-right illustrate the physical significance of the parameter - $K_{diss}$.

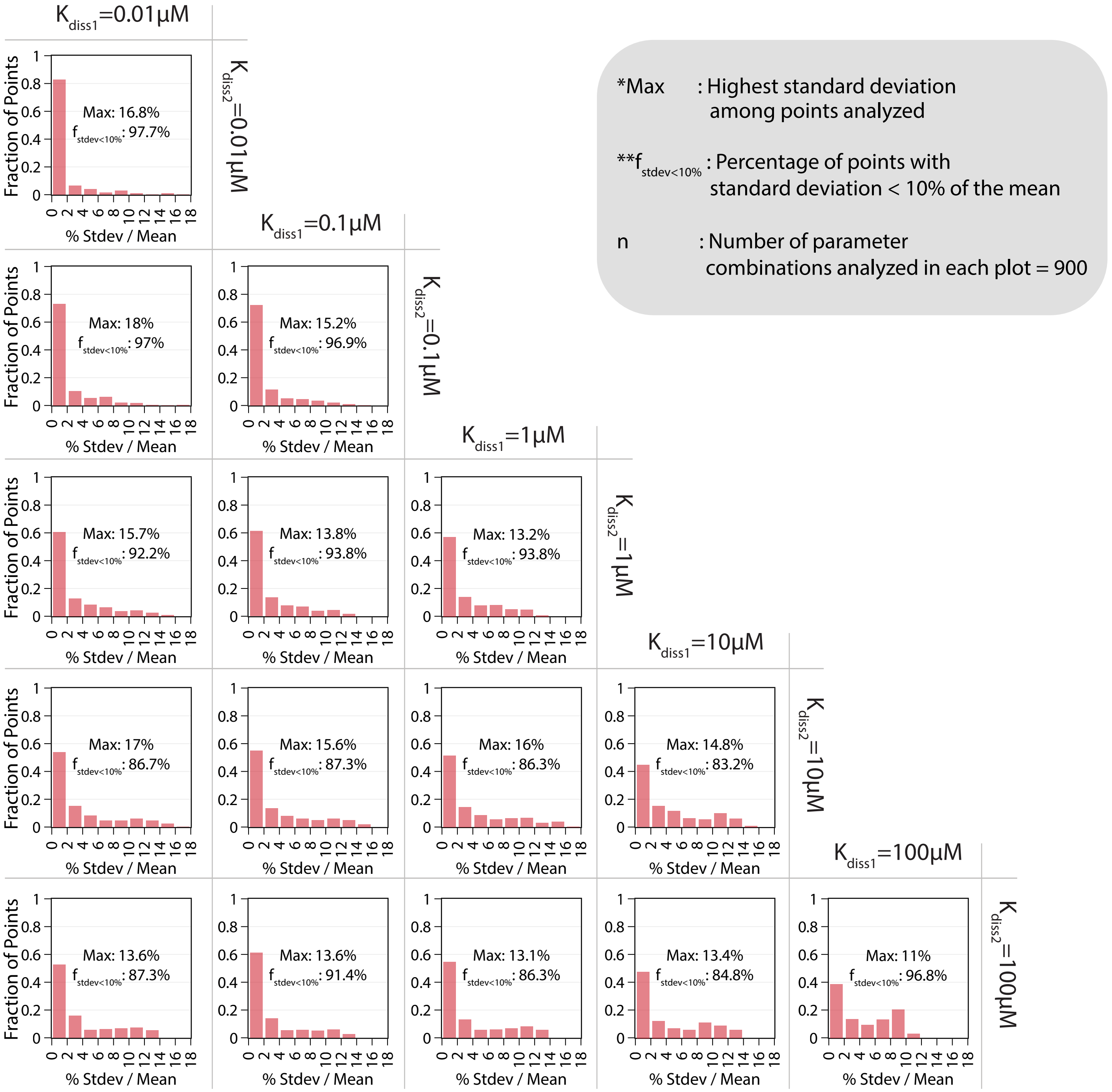


**Figure S5:** The distribution of standard deviations of the switching times for each parameter set, presented as a % of the mean, calculated for all feasible degradation rates, ordered by combinations of $K_{diss}$ values. The maximum value of standard deviation and the fraction of points with standard deviation $<$ 10% of the mean is shown within each plot.

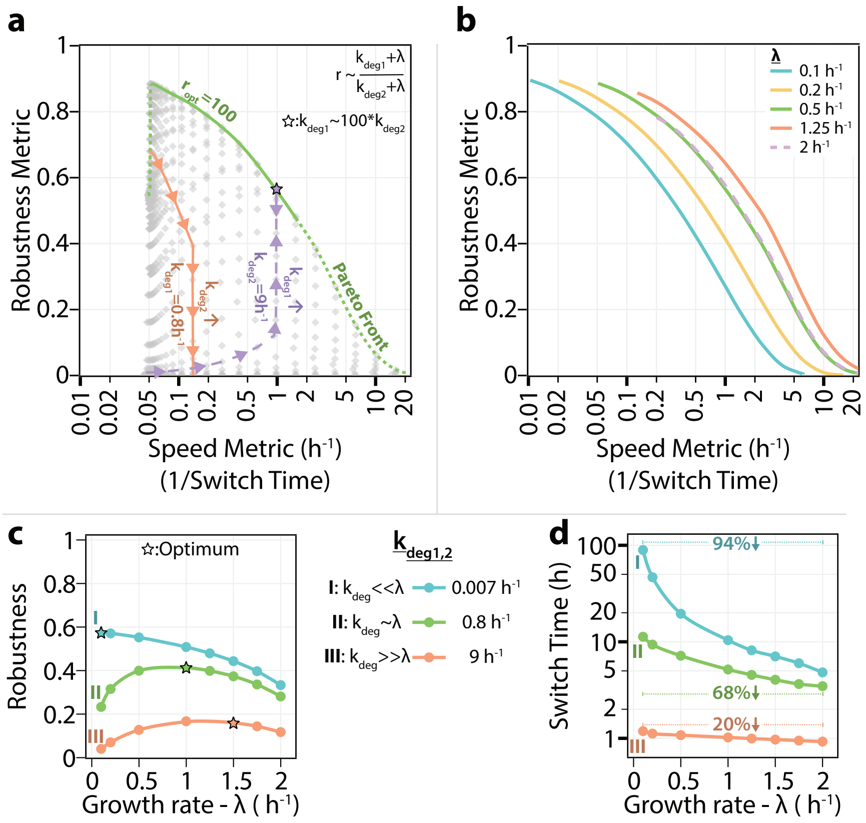


**Figure S6: Growth rate shapes a speed-robustness trade-off and defines optimal switch design. a.** Speed and robustness metrics for all degradation-rate combinations at $K_{diss1} = 0.01$, $K_{diss2} = 1$ µM, with the Pareto front in green. The front is traced when the ratio between the effective degradation rates is roughly equal to 100, i.e. the ratio $r = (k_{deg1} + \lambda)/(k_{deg2} + \lambda)$ equals its optimal value $r_{opt} \sim 100$ (star: $k_{deg1} = 100^{*}k_{deg2}$). The dashed portion of the Pareto front indicates the region where having $r_{opt} \sim 100$ requires degradation rates outside the parameter grid used (i.e. $k_{deg2} < 0.001$ or $k_{deg1} > 1000$). Two trajectories show the effect of tuning one repressor at a time: varying $k_{deg2}$ at fixed $k_{deg1} = 0.8$ h$^{-1}$ (orange) and varying $k_{deg1}$ at fixed $k_{deg2} = 9$ h$^{-1}$ (purple). **b.** Pareto fronts at five growth rates, showing that the achievable frontier itself shifts with growth rate. **c.** Robustness versus growth rate at three representative degradation rates spanning three regimes: (I) $k_{deg} \ll \lambda$ (dilution-dominated), (II) $k_{deg} \sim \lambda$ (mixed), and (III) $k_{deg} \gg \lambda$ (degradation-dominated); stars mark the growth rate of maximal robustness. **d.** Switch time versus growth rate for the same three regimes, with the reduction across the growth-rate range indicated.

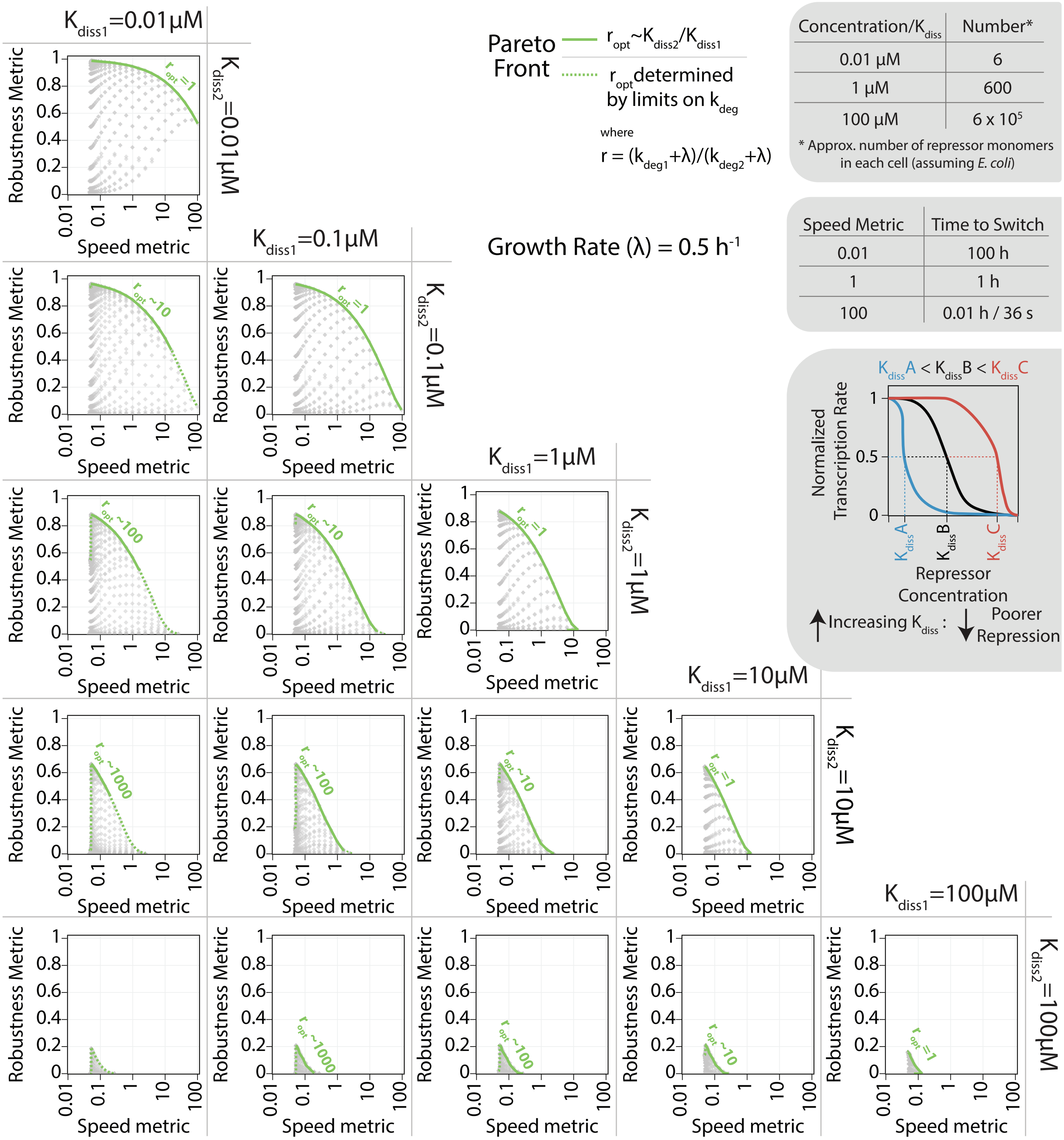


**Figure S7: Pareto front determination from speed - robustness values of all $k_{deg}$ combinations at $\lambda = 0.1h^{-1}$.** The robustness and speed metrics for all combinations of $k_{deg}$ in the simulated parameter grid are plotted for each $K_{diss}$ combination. The pareto front consisting of optimal values of speed-robustness is determined and shown in green for each $K_{diss}$ combination. The ratio $(k_{deg1} + \lambda)/(k_{deg2} + \lambda)$ for points at the pareto front is termed $r_{opt}$ and was found to be equal to the ratio $K_{diss2}/K_{diss1}$ where the corresponding $k_{deg}$ values are within the simulated parameter grid.

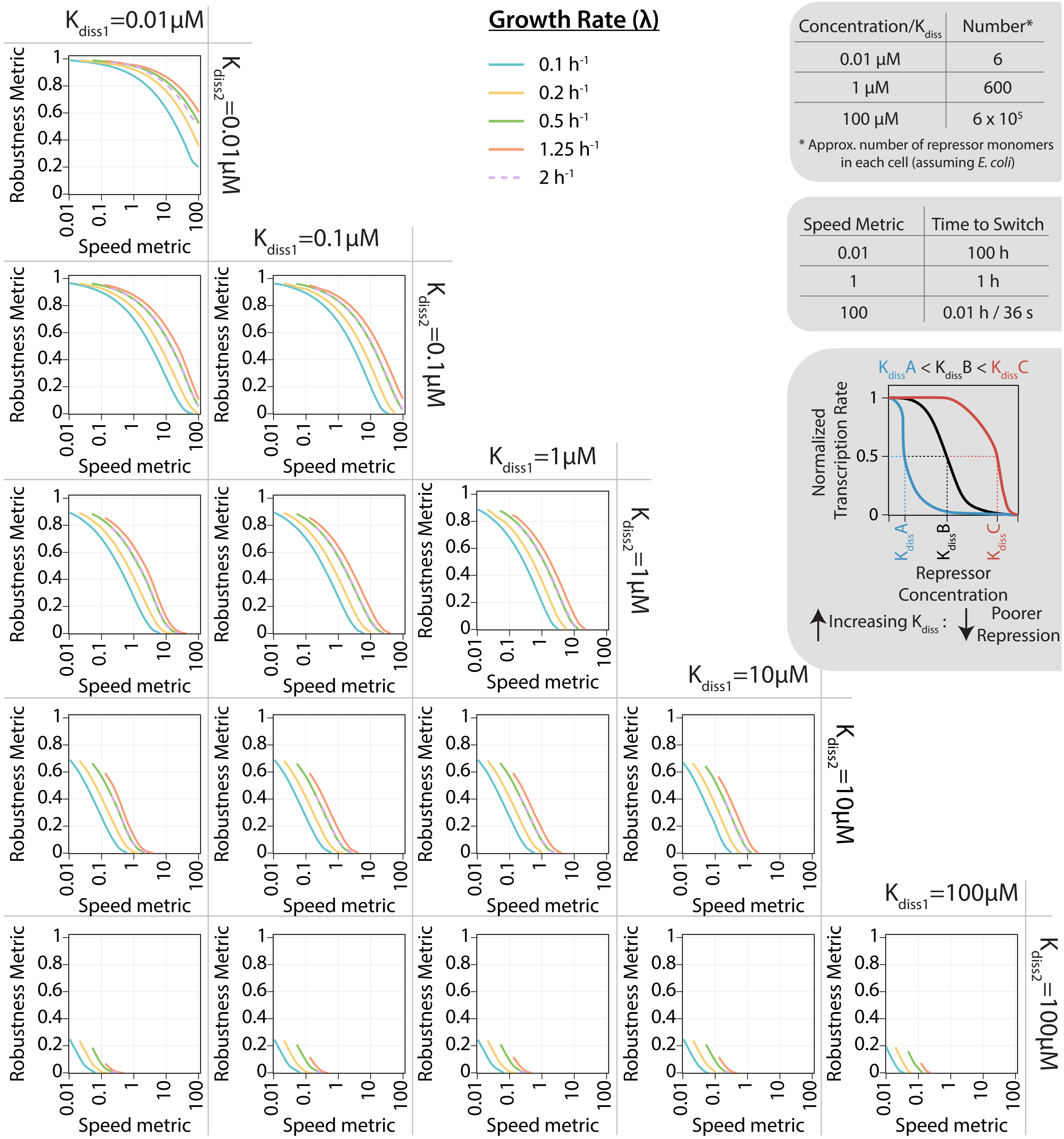


**Figure S8: Pareto front of the speed metric vs robustness metric over entire feasible parametric range.** The relationship between the pareto front of the speed and robustness metrics plotted for all combinations of $K_{diss}$ values that represent the physical limits of experimentally observable values for the parameter computed at five different growth rates ($\lambda$). The shaded boxes to the top-right illustrate the physical significance of the parameter - $K_{diss}$, and the speed metric.

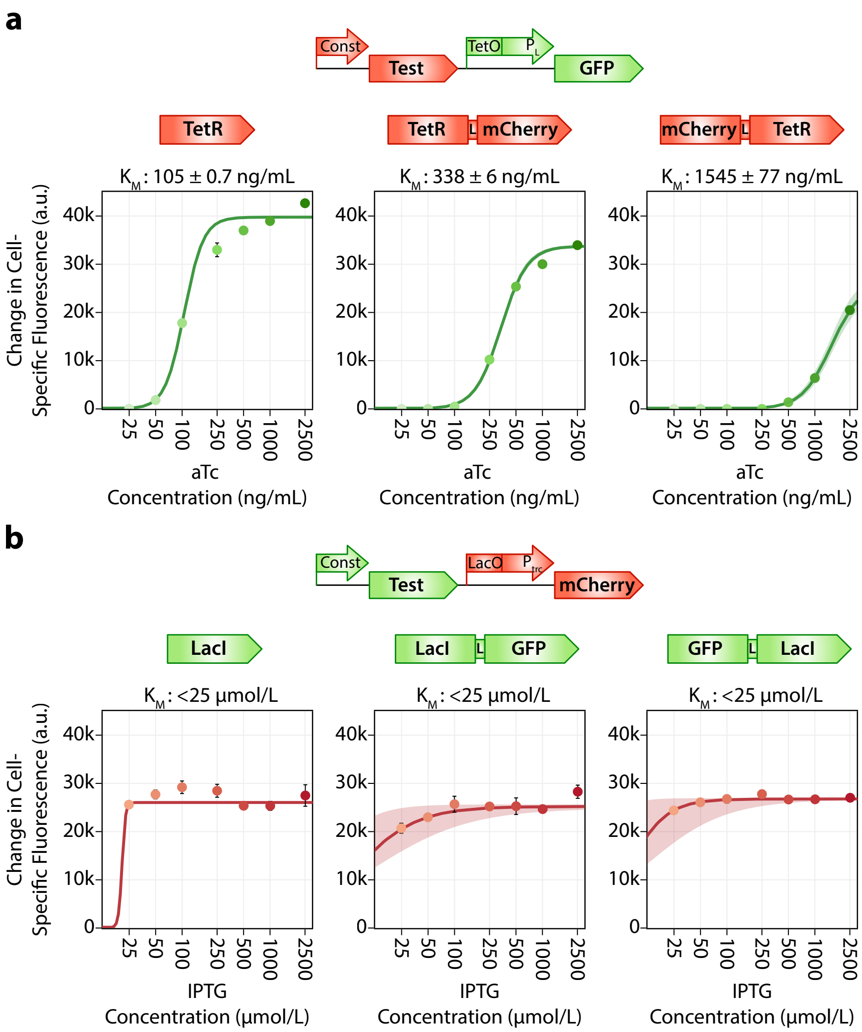


**Figure S9: Test Circuits and Results for Fusion Protein Characterization.** The test circuits to examine the induction of **a.** TetR and **b.** LacI fusion proteins are shown with the corresponding induction profiles for each of the fusion protein variants. 'L' refers to the protein linker used to build the repressor-reporter fusions. For each case, the difference in cell-specific fluorescence between cells induced with various inducer concentrations and uninduced cells is shown. A Hill kinetics equation was fit to this data to obtain the Michaelis-Menten constant $K_M$.

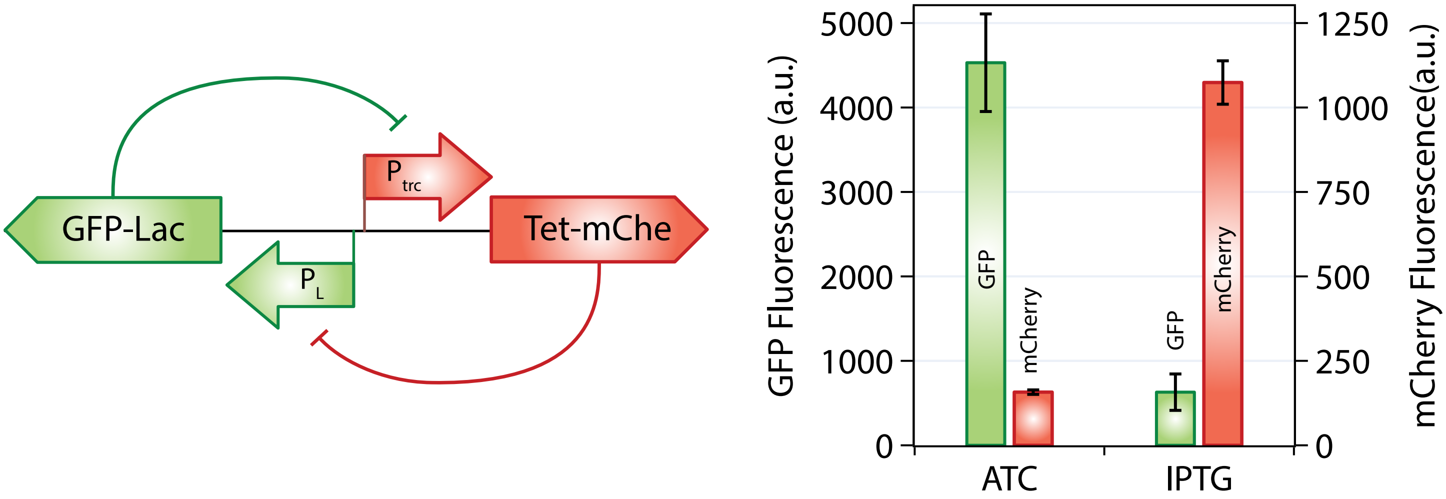


**Figure S10:** Fusion toggle switch variant with the steady state GFP and mCherry fluorescence after growth on media with aTc and IPTG.

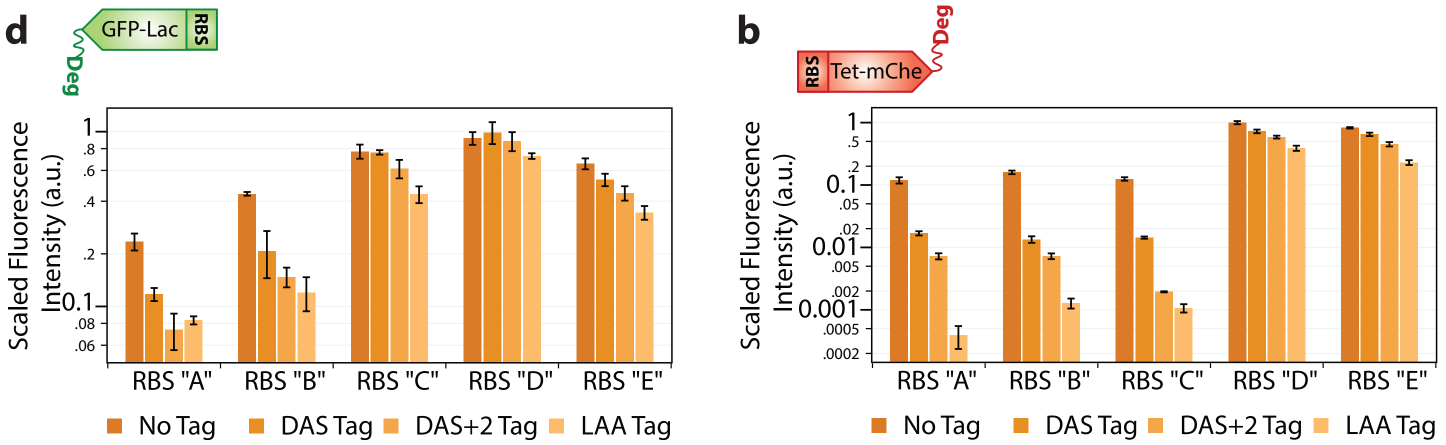


**Figure S11: Characterization of RBS and degradation tag variants of fusion repressor proteins.** Fluorescence intensities of the repressor-reporter fusion constructs determined for each of the 20 RBS-degradation combinatorial variants for **a.**GFP-LacI and **b.** TetR-mCherry, organized by the RBS used. Scaled fluorescence is the fluorescence normalized to the highest value within each plot.

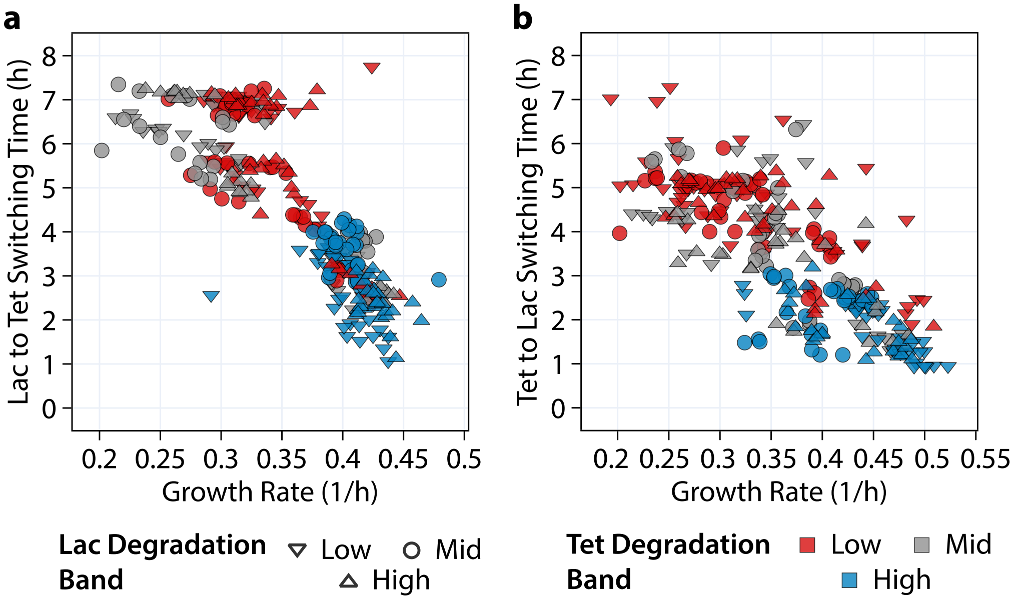


**Figure S12:** Effect of growth rate on switching time. The switching time for a switch from **a.** Lac state to Tet state, and **b.** Tet state to Lac state plotted against the growth rate for individual replicates of all toggle switch constructs classified based on degradation levels estimated in Table S3 and Table S4

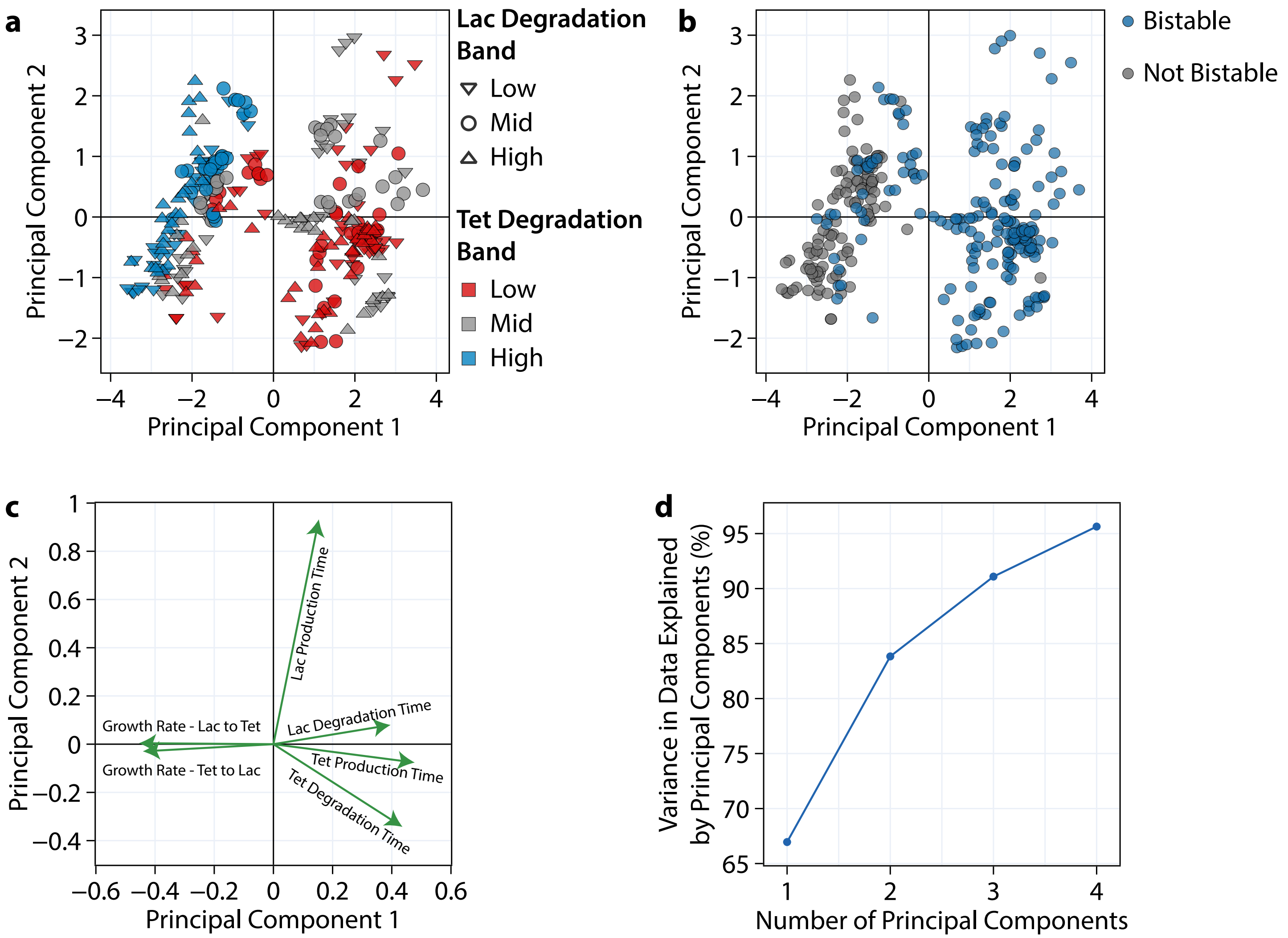


**Figure S13:** A Principal component analysis performed on the various switching times and growth rates of individual replicates of all toggle switch constructs. Scores classified based on **a.** degradation levels estimated in Table S3 and Table S4, b. stability of constructs. **c.** Loadings of each feature. **% Cumulative variance explained by each principal component**

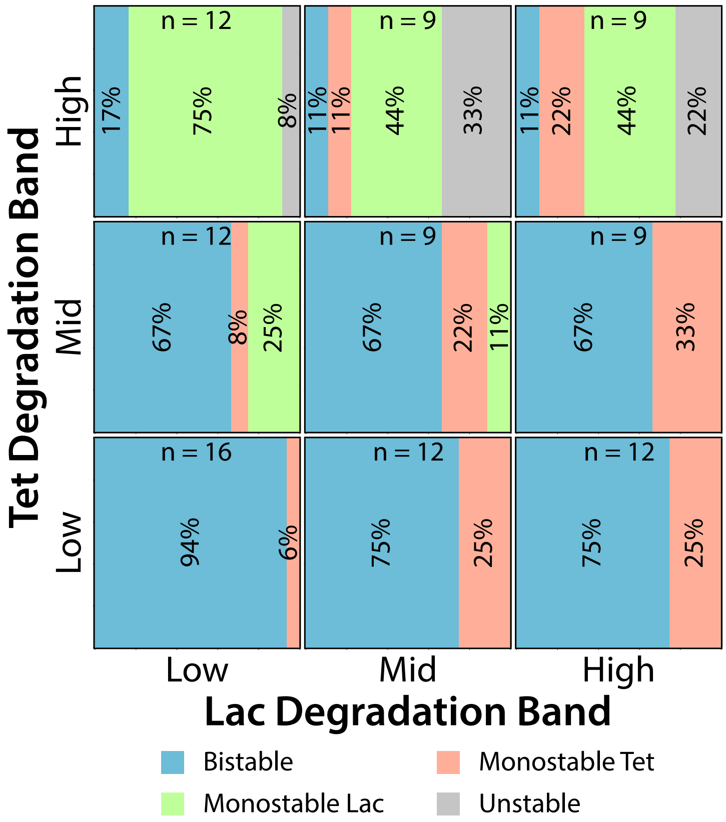


**Figure S14:** Effect of varying degradation levels of the Lac and Tet repressor on the bistability of each toggle switch construct. The percent of bistable, monostable and unstable constructs are shown within the boxes representing each degradation level combination. Constructs were classified into different degradation levels as shown in Table S3 and Table S4. Constructs were classified as bistable if all replicates were bistable.

Figure S14

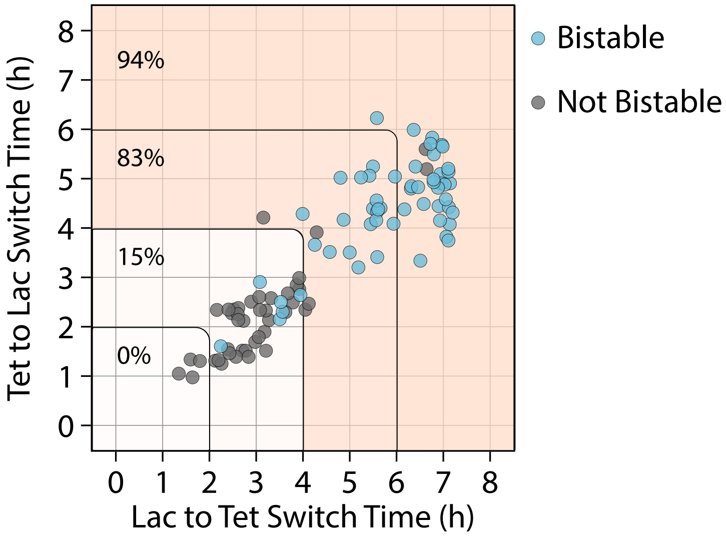


**Figure S15:** The Lac state and Tet state switching times for each construct, classified based on stability. The percent of bistable constructs at each level of Tet and Lac switching times is shown within the plot.

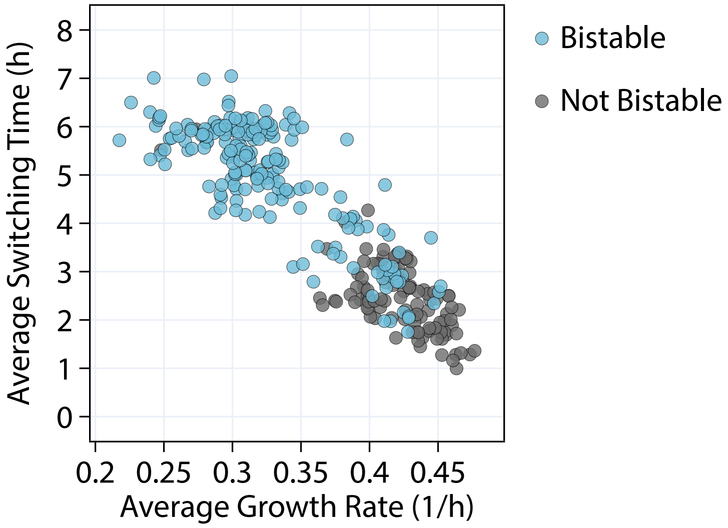


**Figure S16:** A trade-off between the average switching time (calculated as the average of the two state-switching times) and the growth rate of the each strain, classified based on stability for individual replicates.

### S3 Supplementary tables

**Table S1:** Proteome-partitioning parameters used to compute the growth-dependent production rate ceiling $k_{p,max}(\lambda)$.

| Parameter | Description | Value | Source |
|---|---|---|---|
| $r_0$ | Intercept of RNA-to-protein ratio | 0.087 | [4] |
| $k_t$ | Inverse slope of RNA-to-protein ratio | $4.5\,h^{-1}$ | [4] |
| $\rho$ | Ribosomal RNA-to-protein conversion factor | $0.76\,\mu g/\mu g$ | [4] |
| $\phi_{r0}$ | Intercept of ribosomal protein fraction | 0.07 | Derived |
| $w_r$ | Slope of ribosomal protein fraction | $0.17\,h$ | Derived |
| $\phi_{hk}$ | Housekeeping proteome fraction | 0.5 | [4,5] |
| $P_{tot}$ | Total cellular protein concentration | $4\,mM$ | [6] |

**Table S2:** List of ribosome binding sites used in this study

| RBS Name | DNA/RNA Sequence | Description | T.I.R. (RBS calculator v2.1) |
|---|---|---|---|
| pKDL_Tet | CGCCTTCGGCGAAGCTAGGGACGAGAGCTAGC | Reverse Engineered value for RBS of TetR in pKDL071 | 399 |
| pKDL_Lac | ATAGGTCCCAATATAAGGAGAGGGTGTACA | Reverse Engineered value for RBS of LacI in pKDL071 | 8806 |
| pKDL_mCherry | AGTATTAACTATCGTTCAACTGATAGGGAGGCGCCG | Reverse Engineered value for RBS of mCherry in pKDL071 | 575 |
| pKDL_GFP | GCATGCCAGTTGTAGATCAGCATAGCATAGACAACCAAGGAGGTAGTGCAC | Reverse Engineered value for RBS of GFP in pKDL071 | 5627 |
| Fus-Lac | CAGATCCGCTATTCTACAGGGAACCCAATT | Design from query for GFP-LacI with TIR 8800 | 8724 |
| Lac-A | CTGTAAGCTTCCAGTTAAGGAGGTACCGTC | Design from query for GFP-LacI with TIR 10000 | 10764 |
| Lac-B | CTGTAAGCTACCTTTTAAGGAGGTAACGTC | Design from query for GFP-LacI with TIR 25000 | 25027 |

**Table S2 continued from previous page**

| RBS Name | DNA/RNA Sequence | Description | T.I.R. (RBS calculator v2.1) |
|---|---|---|---|
| Lac-C | CTGTATGCTTCCAGTTAA GGAGGTACCGTC | Design from query for GFP-LacI with TIR 50000 | 54985 |
| Lac-D | CTGTATGCTTTCCGTTAA GGAGGTAACGTC | Design from query for GFP-LacI with TIR 100000 | 98299 |
| Lac-E | CTGTATGCTTCCCGTTAA GGAGGTACCGTC | Design from query for GFP-LacI with TIR 150000 | 145449 |
| Fus-Tet | GTCTCATTCCTCGGTTAA AGATCTACTACT | Design from query for TetR-mCherry with TIR 500 | 465 |
| Tet-A | GTCTCGTTAAACGGTTAA CGACGTTATACT | Design from query for TetR-mCherry with TIR 1000 | 1272 |
| Tet-B | GTCTCGTTATTCGGTTAA CGACGGTCTACT | Design from query for TetR-mCherry with TIR 2500 | 2605 |
| Tet-C | GTCTCGTTAAACGGTTAA CGAGGTCATACT | Design from query for TetR-mCherry with TIR 5000 | 5641 |
| Tet-D | GTCTCGTTAAACGGTTAA GGACGGCATACT | Design from query for TetR-mCherry with TIR 10000 | 11590 |
| Tet-E | GTCTCGTTATTCGGTTAA GGACGTTCTACT | Design from query for TetR-mCherry with TIR 15000 | 14845 |

**Table S3:** Degradation rate estimates for TetR constructs based on steady state fluorescence values shown in Figure S11 (Refer to Methods S1.3). Shortlisted variants have been highlighted and classified into three levels of degradation.

| Construct Code | Est Relative Degradation Rate | Degradation Level |
|---|---|---|
| A - * | 1 | |
| A - DAS | 6.79 | |
| A - DAS+2 | 15.58 | |
| A - LAA | 278.44 | |
| B - * | 1 | |
| B - DAS | 11.72 | Mid |
| B - DAS+2 | 21.87 | High |
| B - LAA | 122.81 | |
| C - * | 1 | Low |
| C - DAS | 8.46 | |
| C - DAS+2 | 62.58 | High |
| C - LAA | 114.4 | High |
| D - * | 1 | |
| D - DAS | 1.51 | |
| D - DAS+2 | 1.76 | |
| D - LAA | 2.63 | Mid |
| E - * | 1 | Low |
| E - DAS | 1.33 | Low |
| E - DAS+2 | 2.15 | Low |
| E - LAA | 3.82 | Mid |

**Table S4:** Degradation rate estimates for Lac constructs based on steady state fluorescence values shown in Figure S11 (Refer to Methods S1.3). Shortlisted variants have been highlighted and classified into three levels of degradation.

| Construct Code | Est Relative Degradation Rate | Degradation Level |
|---|---|---|
| A - * | 1 | Low |
| A - DAS | 1.97 | |
| A - DAS+2 | 1.87 | |
| A - LAA | 1.82 | |
| B - * | 1 | Low |
| B - DAS | 2.57 | High |
| B - DAS+2 | 2.14 | |
| B - LAA | 3.32 | |

**Table S4 continued from previous page**

| Construct Code | Est Degradation Rate | Degradation Level |
|---|---|---|
| C - * | 1.02 | Low |
| C - DAS | 1 | |
| C - DAS+2 | 1.1 | |
| C - LAA | 2.25 | High |
| D - * | 1.03 | Low |
| D - DAS | 1 | |
| D - DAS+2 | 1.11 | Mid |
| D - LAA | 1.69 | Mid |
| E - * | 1 | |
| E - DAS | 1.01 | |
| E - DAS+2 | 1.34 | Mid |
| E - LAA | 2.79 | High |

**Table S5:** Degradation tag nucleotide sequences used in toggle switch variants

| Degradation Tag | Repressor | Terminal Nucleotide Sequence |
|---|---|---|
| * | Lac | TGA |
| | Tet | TAA |
| DAS | Lac | GCGGCGAACGATGAAAACTATGCGGATGCGAGCTGA |
| | Tet | GCGGCGAACGATGAAAACTATGCGGATGCGAGCTAA |
| DAS+2 | Lac | GCGGCGAACGATGAAAACTATAACTATGCGGATGCGAGCTGA |
| | Tet | GCGGCGAACGATGAAAACTATAACTATGCGGATGCGAGCTAA |
| LAA | Lac | GCGGCGAACGATGAAAACTATGCGCTGGCGGCGTGA |
| | Tet | GCGGCGAACGATGAAAACTATGCGCTGGCGGCGTAA |

**Table S6:** Description of plasmid sequences attached as supplementary files

| Name | Description |
|---|---|
| pKDL071.gb | Template toggle switch described and made by Litcofsky et al (2012) |
| pTSFusTest - LacI.gb | Plasmid to test repressor effectiveness of LacI |
| pTSFusTest - LacI-GFP.gb | Plasmid to test repressor effectiveness of LacI-GFP fusion protein |
| pTSFusTest - GFP-LacI.gb | Plasmid to test repressor effectiveness of GFP-LacI fusion protein |
| pTSFusTest - TetR.gb | Plasmid to test repressor effectiveness of TetR |
| pTSFusTest - TetR-mCherry.gb | Plasmid to test repressor effectiveness of TetR-mCherry |
| pTSFusTest - mCherry-TetR.gb | Plasmid to test repressor effectiveness of mCherry-TetR |
| pTSFusion.gb | Initial fusion toggle switch construct with TetR-mCherry and GFP-LacI fusion proteins |

**Table S6 continued from previous page**

| Name | Description |
| --- | --- |
| pTSFusion_VariantsTemplate.gb | Template for construction of RBS-degradation tag variants of fusion toggle switch. Nucleotide sequences corresponding to RBS and degradation regions in each variant used in this study can be obtained from Table S2 and Table S5 respectively. Plugging these nucleotide sequences into this template sequence in the annotated RBS and degradation tag regions in this sequence file gives the nucleotide sequences of all plasmids used in this study |

### Bibliography


[1] Jonathan M Monk, Colton J Lloyd, Elizabeth Brunk, Nathan Mih, Anand Sastry, Zachary King, Rikiya Takeuchi, Wataru Nomura, Zhen Zhang, Hirotada Mori, and et al. iml1515, a knowledgebase that computes escherichia coli traits. *Nature Biotechnology*, 35(10):904–908, Oct 2017.

[2] Kaushik Raj, Naveen Venayak, and Radhakrishnan Mahadevan. Novel two-stage processes for optimal chemical production in microbes. *Metab. Eng.*, 62:186–197, nov 2020.

[3] Pauli Virtanen, Ralf Gommers, Travis E. Oliphant, Matt Haberland, Tyler Reddy, David Cournapeau, Evgeni Burovski, Pearu Peterson, Warren Weckesser, Jonathan Bright, Stéfan J. van der Walt, Matthew Brett, Joshua Wilson, K. Jarrod Millman, Nikolay Mayorov, Andrew R.J. Nelson, Eric Jones, Robert Kern, Eric Larson, C. J. Carey, İlhan Polat, Yu Feng, Eric W. Moore, Jake VanderPlas, Denis Laxalde, Josef Perktold, Robert Cimrman, Ian Henriksen, E. A. Quintero, Charles R. Harris, Anne M. Archibald, Antônio H. Ribeiro, Fabian Pedregosa, Paul van Mulbregt, Aditya Vijaykumar, Alessandro Pietro Bardelli, Alex Rothberg, Andreas Hilboll, Andreas Kloeckner, Anthony Scopatz, Antony Lee, Ariel Rokem, C. Nathan Woods, Chad Fulton, Charles Masson, Christian Häggström, Clark Fitzgerald, David A. Nicholson, David R. Hagen, Dmitrii V. Pasechnik, Emanuele Olivetti, Eric Martin, Eric Wieser, Fabrice Silva, Felix Lenders, Florian Wilhelm, G. Young, Gavin A. Price, Gert Ludwig Ingold, Gregory E. Allen, Gregory R. Lee, Hervé Audren, Irvin Probst, Jörg P. Dietrich, Jacob Silterra, James T. Webber, Janko Slavič, Joel Nothman, Johannes Buchner, Johannes Kulick, Johannes L. Schönberger, José Vinícius de Miranda Cardoso, Joscha Reimer, Joseph Harrington, Juan Luis Cano Rodríguez, Juan Nunez-Iglesias, Justin Kuczynski, Kevin Tritz, Martin Thoma, Matthew Newville, Matthias Kümmerer, Maximilian Bolingbroke, Michael Tartre, Mikhail Pak, Nathaniel J. Smith, Nikolai Nowaczyk, Nikolay Shebanov, Oleksandr Pavlyk, Per A. Brodtkorb, Perry Lee, Robert T. McGibbon, Roman Feldbauer, Sam Lewis, Sam Tygier, Scott Sievert, Sebastiano Vigna, Stefan Peterson, Surhud More, Tadeusz Pudlik, Takuya Oshima, Thomas J. Pingel, Thomas P. Robitaille, Thomas Spura, Thouis R. Jones, Tim Cera, Tim Leslie, Tiziano Zito, Tom Krauss, Utkarsh Upadhyay, Yaroslav O. Halchenko, and Yoshiki Vázquez-Baeza. SciPy 1.0: fundamental algorithms for scientific computing in Python. *Nat. Methods*, 17(3):261–272, 2020.

[4] Matthew Scott, Carl W. Gunderson, Eduard M. Mateescu, Zhongge Zhang, and Terence Hwa. Interdependence of cell growth and gene expression: Origins and consequences. *Science*, 330(6007):1099–1102, Nov 2010.

[5] Matteo Mori, Terence Hwa, Olivier C. Martin, Andrea De Martino, and Enzo Marinari. Constrained allocation flux balance analysis. *PLOS Computational Biology*, 12(6), Jun 2016.

[6] F.C. Neidhardt. *Escherichia Coli and Salmonella: Cellular and Molecular Biology.* Number v. 1-2 in Escherichia Coli and Salmonella: Cellular and Molecular Biology. ASM Press, 1996.
